# Biomimetic S-Adenosylmethionine Regeneration Starting from Different Byproducts Enables Biocatalytic Alkylation with Radical SAM Enzymes

**DOI:** 10.1101/2022.09.26.509380

**Authors:** Lukas Gericke, Dipali Mhaindarkar, Lukas Karst, Sören Jahn, Marco Kuge, Michael K. F. Mohr, Jana Gagsteiger, Nicolas V. Cornelissen, Xiaojin Wen, Silja Mordhorst, Henning J. Jessen, Andrea Rentmeister, Florian P. Seebeck, Gunhild Layer, Christoph Loenarz, Jennifer N. Andexer

## Abstract

*S*-Adenosylmethionine (SAM) is an enzyme cofactor involved in methylation, aminopropyl transfer, and radical reactions. This versatility renders SAM-dependent enzymes of great interest in biocatalysis. The usage of SAM analogues adds to this diversity. However, high cost and instability of the cofactor impedes the investigation and usage of these enzymes. While SAM regeneration protocols from the methyltransferase (MT) byproduct *S*-adenosylhomocysteine are available, aminopropyl transferases and radical SAM enzymes are not covered. Here, we report an efficient one-pot system to supply or regenerate SAM and SAM analogues for all three enzyme classes. The system’s flexibility is showcased by the transfer of an ethyl group with a cobalamin-dependent radical SAM MT using *S*-adenosylethionine as a cofactor. This shows the potential of SAM (analogue) supply and regeneration for the application of diverse chemistry, as well as for mechanistic studies using cofactor analogues.

## Introduction

*S-*adenosylmethionine (SAM, AdoMet) is one of the most abundant and versatile enzyme cofactors. The sulfonium ion is widely known as one of nature’s methyl group donors and for its role in the biosynthesis of a wide range of natural products.^[1–3]^ Methylation is essential for epigenetic regulation *via* nucleobase and histone methylation and consequently closely connected to human health and disease.^[4,5]^ Especially in drug synthesis, methylation is a highly effective modification that can increase the potency of a substance up to three orders of magnitude, leading to the term “magic methyl effect”.^[6,7]^ SAM-dependent enzymes have the potential to provide an environmentally friendly alternative to commonly used toxic, and potentially unselective alkylating agents.^[8]^

Conventional SAM-dependent methyltransferases (MTs, E.C. 2.1.1) use SAM as a methyl donor with a nucleophilic substrate in an S_N_2 reaction, forming *S*-adenosylhomocysteine (SAH) as a byproduct (Figure 1a).^[1–3]^ The biosynthesis of many pharmacologically active natural products such as galantamine and caffeine involves MTs (Figure 1b). In addition, other SAM-dependent enzymes perform versatile reactions while using different substructures of the cofactor.^[2,3,9]^ Aminocarboxypropyl transferases (ACPT, e. g. E.C. 2.5.1.38) transfer the amino-carboxypropyl moiety of SAM to various substrates.^[10–12]^ After decarboxylation of SAM, the aminopropyl side chain provides building blocks for the biosynthesis of polyamines by aminopropyl transferases (APT, e. g. E.C. 2.5.1.16);^[13]^ in both cases, 5′-methylthioadenosine (MTA) is formed as the SAM-derived byproduct (Figure 1a). Simple linear polyamines such as spermidine and spermine occur in all forms of life; they are important for diverse cellular processes, including apoptosis, cell differentiation, gene expression, homeostasis, and signalling.^[13,14]^ Polyamines are also used as building blocks for polyamine alkaloids with multiple biological functions.^[15]^ This group of natural products contains acylated and alkylated polyamines, as well as compounds containing other building blocks including terpenes and nucleobases. Examples are the cytotoxic compounds motuporanine G-I and the spider toxin PA3334G (Figure 1c).^[15,16]^

**Figure 1.**
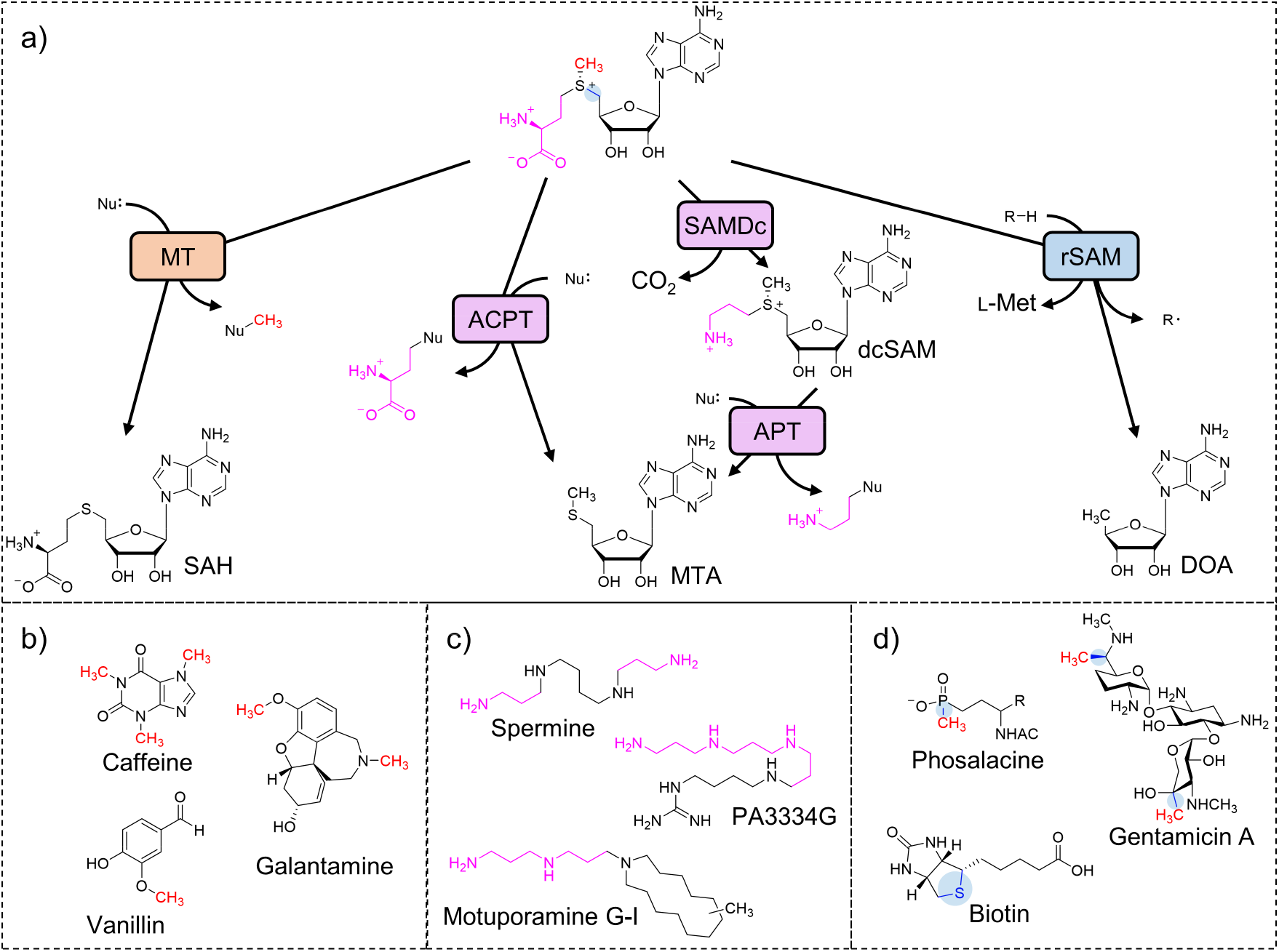
SAM is a highly versatile cofactor involved in a wide range of reactions. **a)** SAM-dependent enzymes can be divided into three large groups based on the structural moieties transferred: methyl transferases (MT, red), enzymes transferring the aminocarboxypropyl moiety (ACPT) or the aminopropyl moiety (APT) after decarboxylation by SAM decarboxylase (SAMDc) (pink), and radical SAM enzymes (rSAM, blue). The substrate radical is displayed as a common intermediate for the diverse reactions catalysed by radical SAM enzymes. **b)** Examples for methylated drugs and food additives. **c)** Examples for polyamines. **d)** Examples for radical SAM enzyme products.

The discovery of the radical SAM superfamily (PF04055) dramatically increased the scope of SAM-dependent reactions and consequently potential applications. The members of this superfamily use a SAM-derived 5’-deoxyadenosyl radical (DOA•) to initiate radical chemistry for a broad range of synthetically challenging reactions (Figure 1a).^[17]^ These include methylations at unactivated carbons, other C-C bond formations as well as rearrangements such as ring contractions.^[18–21]^ Radical SAM enzymes are present in the biosynthetic gene clusters of countless natural products, including biotin and the antibiotics gentamicin and phosalacine (Figure 1d).^[18,22]^ Alternatively to abstracting a hydrogen atom from the substrate [resulting in 5′-deoxyadenosine (DOA) as a byproduct], the DOA• radical can be added to an alkene resulting in an adenosylation.^[23]^ Regarding methylation reactions, particularly class B radical SAM MTs have gained interest due to their ability to catalyse challenging methylations on sp^3^-hybridised C or P atoms.^[24–29]^ In addition to the SAM molecule used to generate the DOA• radical, class B radical SAM MTs utilise a secondSAM molecule to methylate cobalamin, which serves as a methyl shuttle.^[30]^

The diversity of SAM-dependent reactions renders them of great interest for application in organic synthesis and biotechnological processes.^[8,23,31,32]^ In recent years, the large reaction scope of SAM-dependent enzymes has been further expanded by the usage of SAM analogues. MTs have been used together with alkyl analogues of SAM to transfer bulkier residues.^[33–35]^ Radical SAM enzymes have been shown to support the cleavage of *S*-adenosylethionine (SAE) and *S*-adenosyl-L-allyl homocysteine as well as the transfer of various nucleosides to substrate analogues.^[36–38]^

In addition to the high cost and low stability of the SAM cofactor,^[39]^ in many cases SAM-derived byproducts have an inhibitory effect on the corresponding enzymes.^[18,40,41]^ This, and the desire to use methionine/SAM analogues, encouraged the development of multi-enzyme systems for SAM supply or regeneration. Biomimetic SAM supply starts from L-methionine (L-Met) and 5’-adenosine triphosphate (ATP) using methionine adenosyltransferase (MAT, E.C. 2.5.1.6), nature’s SAM-synthesising enzyme.^[42,43]^ Alternatively, chlorinases (E.C. 2.5.1.94) are employed to synthesise SAM from L-Met and 5′-chloro-5′-deoxyadenosine (CldA) in their reverse reaction.^[44]^ Both options are suitable for producing SAM analogues from methionine analogues for the transfer of alternative alkyl groups.^[33,45–47]^ The inhibiting SAM-derived byproduct SAH (in the case of conventional MTs) can be removed using natural salvage enzymes: either SAH hydrolase (SAHH, E.C. 3.3.1.1),^[48]^ or MTA/SAH nucleosidase (MTAN, E.C. 3.2.2.16) yielding adenosine or adenine, respectively. MTAN offers the advantages of its broader substrate range and catalysing an irreversible reaction. In biology, MTAN also cleaves the SAM-derived byproducts DOA (from radical SAM enzymes) and MTA (from ACP transfers and polyamine biosynthesis), yielding the corresponding ribose derivative and adenine.^[40,49]^ SAM supply cascades are well established and have been used for many different scenarios;^[34,46,50–55]^ mainly for conventional SAM-dependent MTs, using the physiological cofactor or SAM analogues.^[33,52,54]^ Radical SAM enzymes have been used in combination with MAT or MTAN for the *in situ* production of cofactor (analogues) or the removal of DOA, respectively.^[56–59]^ Nevertheless, depending on the application, a cofactor regeneration system is desired that does not require stoichiometric amounts of the precursors ATP or CldA. For instance, the low amount of cofactor present can simplify downstream processing of the main product.^[60]^ For conventional SAM-dependent MTs a biomimetic cycle using SAHH has been implemented: the resulting adenosine is rephosphorylated to ATP using polyphosphate (polyP) as a phosphate donor. Optionally, the second SAHH product, L-homocysteine, can be remethylated using homocysteine *S*-methyltransferase (E.C. 2.1.1.10) in combination with methylmethionine as a methyl donor, reaching up to 200 total turnover numbers (TTN).^[61,62]^ This system is also applicable for L-Met and nucleobase derivatives. Another approach for SAM regeneration uses the reverse reaction of a halide MT (HMT, E.C. 2.1.1.165) to directly remethylate SAH using methyliodide with up to 500 TTN;^[63]^ this has been expanded to the use of other alkyl halides for general alkylation.^[64–67]^ Both regeneration systems have in common that they rely on SAH formed in the MT reaction. Hence, they exclusively support MTs since APTs and radical SAM enzymes produce MTA and DOA, respectively. During the development of both systems adenine was identified as a dead-end byproduct that cannot be regenerated in the implemented setups and is consequently lost as a cofactor building block. In the HMT system, this undesired activity was traced back to the co-purified MTAN enzyme from *E. coli*, and could be resolved by using an *mtan*-knockout *E. coli* strain for heterologous expression.^[63]^ For the biomimetic multi-step system this was also partially successful, although a small amount of adenine is additionally formed through a nucleosidase side activity observed for SAHH from *Mus musculus*.^[62]^

In this work, we used adenine as a starting point for an *in vitro* biomimetic SAM regeneration cycle applicable for all three SAM-derived byproducts: SAH, DOA, and MTA. Such a flexible system has the potential to unlock polyamine synthesis and the vast chemistry of radical SAM enzymes for SAM regeneration, and the support of SAM analogues for product diversification and as a tool for functional characterisation. In addition, we were interested to test radical SAM MTs for their ability to transfer larger alkyl chains, as it is established for non-radical MTs (Figure 2).

**Figure 2.**
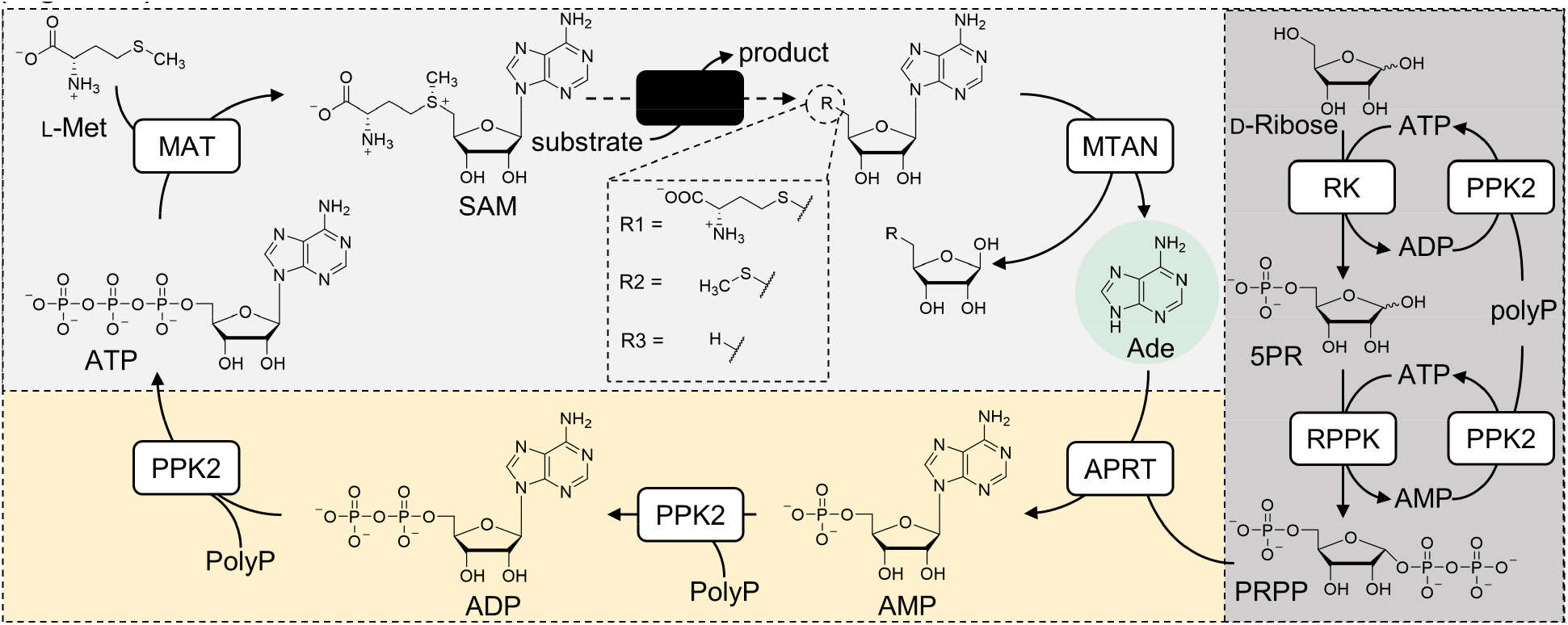
The SAM regeneration system supporting MTs, APTs and radical SAM enzymes envisioned in this work. The system is based on a SAM supply cascade consisting of MAT for SAM supply from L-Met or an analogue, and ATP, SAM-dependent enzyme(s) (black rounded box) and MTAN for byproduct cleavage (light grey). The 5’-residue of the ribosyl moiety depends on the class of SAM-dependent enzymes: R1 for MTs, R2 for APTs and R3 for radical SAM enzymes. The adenine (Ade) produced in the MTAN reaction is the starting point for SAM regeneration by condensing it with 5-phosphoribosyl-1-pyrophosphate (PRPP) synthesised by a ribose phosphorylation cascade (dark grey). The AMP formed is phosphorylated twice to yield ATP, completing the cycle (yellow).

## Results and Discussion

### PolyP-based Production of ATP from Adenine

In the natural salvage pathway of nucleotide biosynthesis, ribophosphorylation of adenine is carried out by adenine phosphoribosyltransferases (APRT, E.C. 2.4.2.7) using 5-phosphoribosyl-1-pyrophosphate (PRPP) as a cofactor. ^[68]^ We aimed for a biomimetic *in situ* synthesis of PRPP from D-ribose using ribokinase (RK, E.C. 2.7.1.15) and D-ribose-5-phosphate pyrophosphokinase (RPPK, E.C. 2.7.6.1) (Figure 2, dark grey box, Figure S1). This approach has been successfully used before as one-pot reactions to produce different types of nucleotides from their corresponding nucleobase and sugars.^[69–73]^ Both, RK and RPPK, use ATP for the phosphorylation reactions. Different regeneration systems have been used for the rephosphorylation of ADP (RK reaction) or AMP (RPPK reaction). Based on our prior experience, we chose a set of PPK2 enzymes established in our laboratory: *Aj*PPK2 from *Acinetobacter johnsonii* and *Sm*PPK2 from *Sinorhizobium meliloti* for the polyP-dependent phosphorylation of AMP and ADP, respectively (details in Figure S2).^[61]^

This polyP-powered adenine-to-ATP module can now be universally linked with the linear SAM supply/SAM-derived byproduct degradation cascades (Figure 2, light grey box) for all three types of SAM-dependent enzymes in a one-pot system. The performance of each enzyme class (conventional MTs, radical SAM enzymes, and APTs) was first tested in the SAM supply cascade containing MAT and MTAN (Figure 2, light grey box) and then integrated into the cycle (Figure 2).

### SAM Regeneration from SAH via Adenine/MTAN Works with Conventional MTs Using an S_N_2 Mechanism

The incorporation of MTs into the system was tested with anthranilate *N*-MT from *Ruta graveolens* (*Rg*ANMT, E.C. 2.1.1.111) and catechol *O*-MT from *Rattus norvegicus* (*Rn*COMT, E.C. 2.1.1.6) (Figure 3). These enzymes have been successfully integrated into the linear SAM supply cascade containing MAT and MTAN, as well as the biomimetic SAHH cycle, and are thus a well-suited comparison.^[55,61,62]^ In the linear cascade, *Rn*COMT was investigated for its byproduct inhibition, showing the benefits of MTAN addition.^[55]^ The combined regeneration system contained the purified enzymes (Figure S1), starting materials, and salts and was incubated for 20 h. Samples were analysed by HPLC (details in the experimental section). Three different AMP concentrations were tested for each enzyme, corresponding to 10×, 50× and 100× excess of the substrate (2 mM) over AMP. As a parameter for the performance of the system, the number of catalytic cycles was determined as the total turnover number (TTN) by the ratio of product formed to added cofactor after 20 h.^[74,75]^

**Figure 3.**
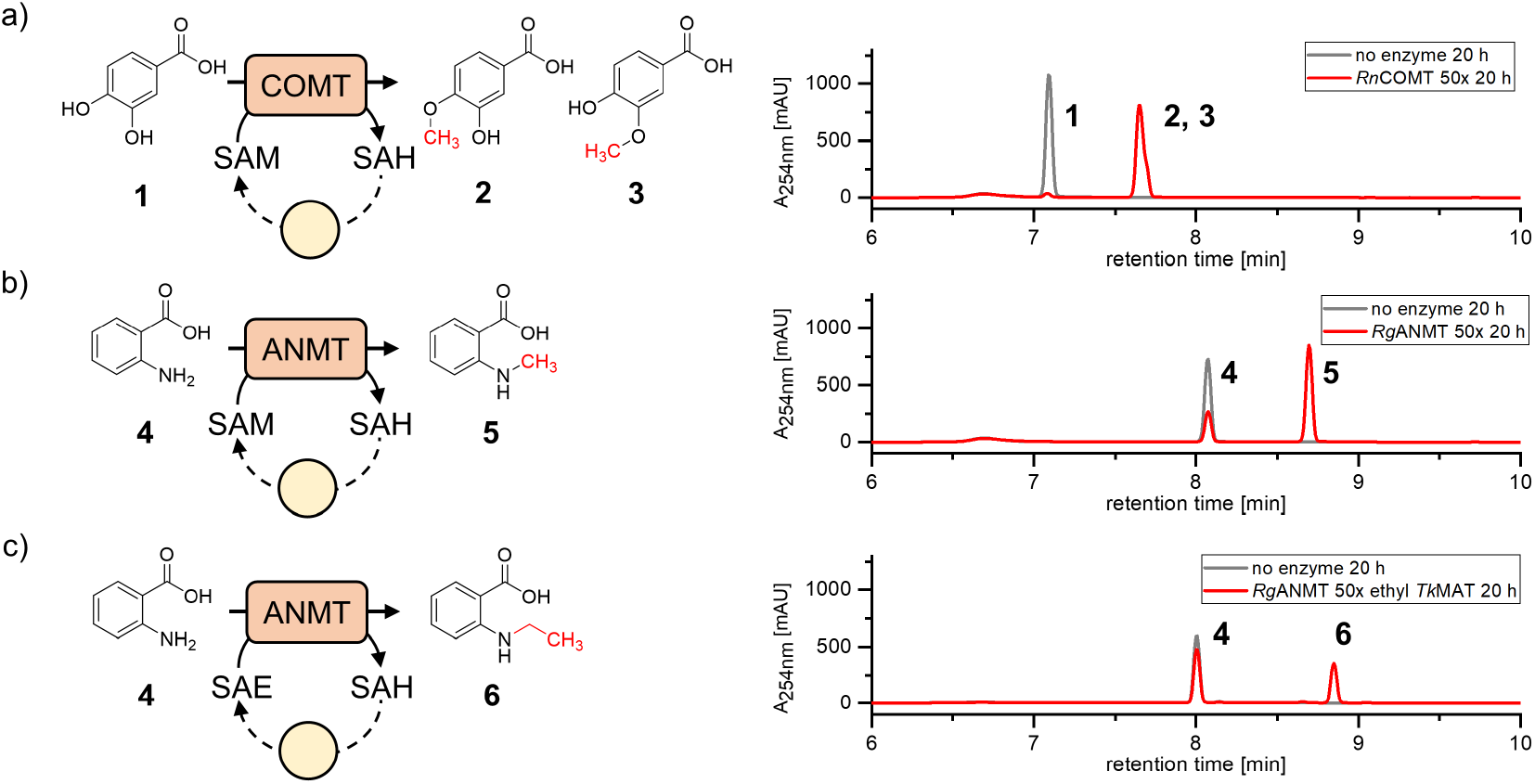
Reactions and HPLC-analysis of conventional MTs using the SAM regeneration *via* adenine (yellow circle). **a)** 3,4-Dihydroxybenzoic acid (**1**) was methylated to yield a mixture of isovanillic acid (**2**) and vanillic acid (**3**). The chromatogram shows conversion of **1** to a mixture of **2** and **3** after 20 h in experiments containing 50× excess of substrate over AMP compared to a no enzyme control containing (**2** and **3** were not separated by the HPLC method used). **b)** Anthranilic acid (**4**) was converted to *N*-methylanthranilic acid (**5**). Chromatograms are shown for experiments with 50× excess of substrate over AMP with L-Met present. The Assays showed conversion of **4** to **5**. c) In assays investigating ethylation with the system, L-Eth replaced L-Met and *Tk*MAT replaced *Ec*MAT to support efficient SAE supply. Conversion of **4** to *N*-ethylanthranilic (**6**) was observed.

A 10× excess of substrate over AMP led to full conversion and a TTN of 10 with both MTs. A TTN of 46 ± 4 was determined for *Rn*COMT with 50× excess of substrate (Figure 3a); as observed before,^[62]^ *Rg*ANMT was less efficient, nevertheless with a conversion of 65% (TTN 33 ± 1, Figure 3b). The highest TTN measured for this system was 80 ± 5 for *Rn*COMT at 100× substrate excess over AMP (Table 1, Figure S3a).

**Table 1.** Combined quantitative data for the SAM regeneration system *via* MTAN/Ade.

| Enzyme | c(AMP) [mM] | substrate excess | conversion [%] | TTN |
| --- | --- | --- | --- | --- |
| *RnCOMT* | 0.2 | 10× | 100 ± 0 | 10 ± 0 |
|  | 0.04 | 50× | 93 ± 9 | 46 ± 4 |
|  | 0.02 | 100× | 80 ± 5 | 80 ± 5 |
| *RgANMT* | 0.2 | 10× | 97 ± 3 | 10 ± 0 |
|  | 0.04 | 50× | 65 ± 1 | 33 ± 1 |
|  | 0.02 | 100× | 67 ± 12 | 67 ± 12 |
| *EcSpeS* | 0.2 | 10× | 96 ± 1 | 10 ± 0 |
|  | 0.04 | 50× | 82 ± 4 | 41 ± 2 |
|  | 0.02 | 100× | 57 ± 10 | 57 ± 10 |
| *AtSpeS* | 0.2 | 10× | 98 ± 0 | 10 ± 0 |
|  | 0.04 | 50× | 79 ± 8 | 40 ± 4 |
|  | 0.02 | 100× | 55 ± 11 | 55 ± 11 |
| QCMT | 0.04 | 10× | 90 ± 3 | 18 ± 1 |
|  | 0.02 | 25× | 56 ± 22 | 28 ± 11 |

To investigate the ability of the SAM regeneration system to support alkylation reactions other than methylations, MAT from *Thermococcus kodakarensis* (*Tk*MAT) was used for SAE generation as the enzyme showed favourable properties towards the acceptance of SAM analogues.^[76,77]^ L-Ethionine (L-Eth) replaced L-Met in these experiments. *Rg*ANMT was used for ethyl transfer to anthranilic acid for comparison to previous multi enzyme SAM regeneration systems.^[61,62]^ With 50× excess of substrate over AMP, 30% conversion and 15 TTN were observed (Figure 3b), slightly higher than the previously reported 20% for the adenosine/SAHH-based regeneration system.^[61]^ This supports the assumption that MAT and/or ANMT require higher amounts of cofactor building blocks when L-Eth is used, especially as the corresponding linear cascade showed >75% conversion.^[61,62]^

### The SAM Regeneration System Supports Class B Radical SAM MTs by SAM Regeneration from SAH and DOA

From the vast group of radical SAM enzymes, we were especially interested in radical SAM MTs, as these enzymes, in combination with SAM analogues, would dramatically expand the range of biocatalytical alkylation reactions. The cobalamin-dependent glutamine *C*-MT (QCMT) from *Methanoculleus thermophilus* served as a model enzyme. This recently described enzyme is involved in the posttranslational modification of methyl-coenzyme M reductase catalysing the methylation of the sp^3^-C2 of Q418.^[56]^ This enzyme requires two SAM molecules per catalytic cycle: one is homolytically cleaved into L-Met and a DOA• radical, the other serves as a methyl group donor to Co(I) of bound cobalamin to generate methylcobalamin [MeCbl(III)] under release of SAH. The DOA• radical then abstracts a hydrogen atom from C2 of the glutamine residue, resulting in DOA and the corresponding peptidyl radical. The methylation of the peptidyl radical proceeds *via* homolytic cleavage of the cobalt-carbon bond producing the methylated peptide and cob(II)alamin [Cbl(II), Figure 4a].^[56]^ As both, DOA and SAH, are substrates for MTAN^[40]^, the regeneration system should be capable of cleaving both byproducts (SAH and DOA) and regenerating two equivalents of SAM from adenine per methylation reaction.

**Figure 4.**
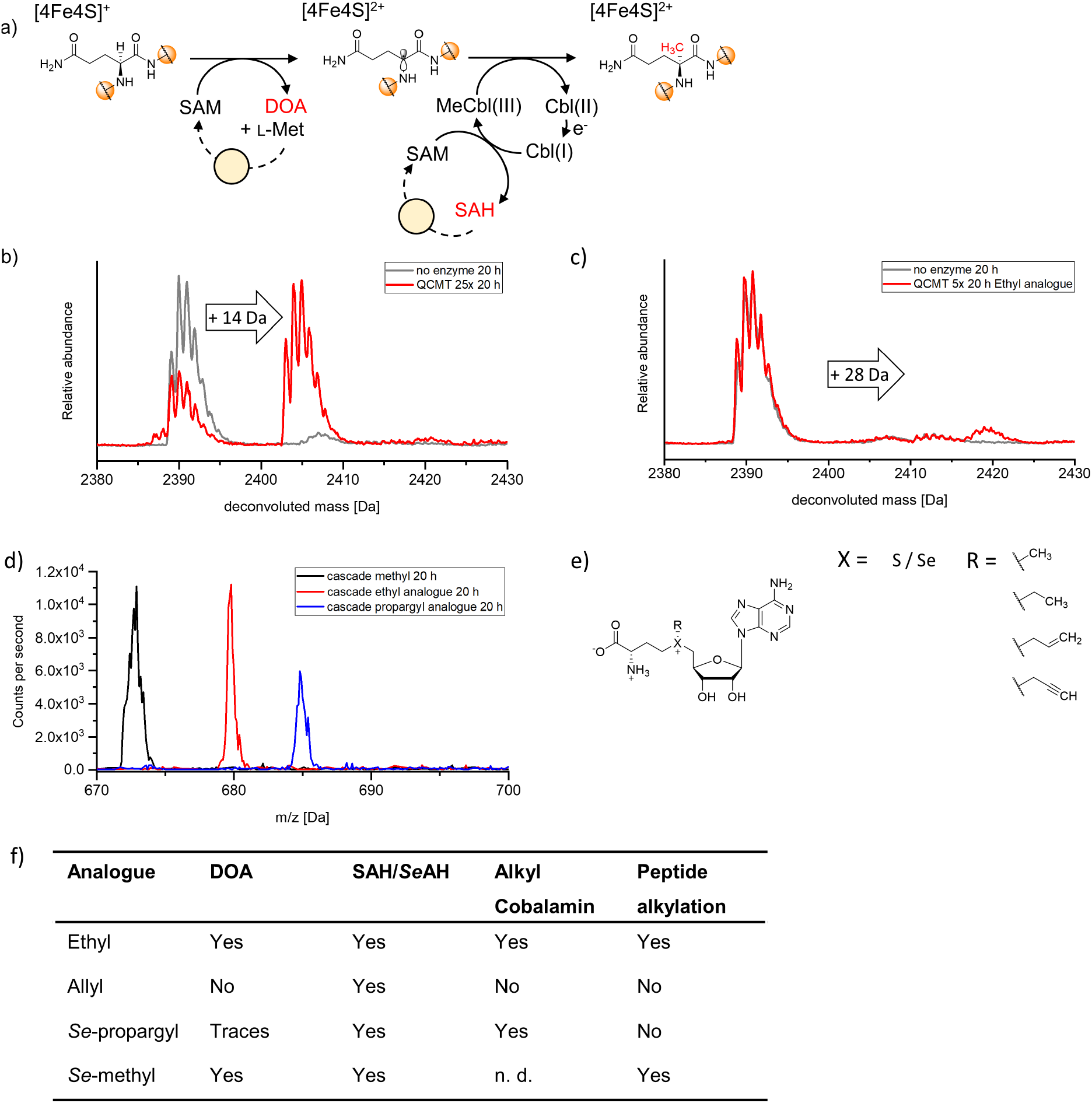
SAM regeneration for class B radical SAM MTs. a) QCMT methylates C2 of a glutamine residue within a peptide chain (orange) under retention of stereochemistry.^[56]^ Two SAM molecules are utilised, one for radical generation, the other for cobalamin methylation. DOA and SAH are formed as byproducts, both can be regenerated to SAM (yellow circle). b) LC-MS analysis of assays with 25× excess of peptide substrate (m = 2390.1 Da) over AMP (50 theoretical TTN) showed a mass shift of +14 Da (m = 2404.1 Da), consistent with methylation. c) Usage of L-Eth instead of L-Met in the SAM regeneration system with 5× excess of peptide substrate over AMP leads to ethylation of the peptide substrate, indicated by a mass shift of +28 Da (m = 2419.0 Da). d) Analysis of cobalamin bound to the enzyme shows evidence for methyl-(m/z 672.9 Da), ethyl-(m/z 679.8 Da) and propargylcobalamin (m/z 684.8 Da) in cascade assays with the respective SAM analogue present. Each cobalamin species was detected as m + 2 H^+^. e) Analogues of SAM tested with QCMT. f) Overview of the observed species in QCMT assays using the individual L-Met analogues.

Initially, the three-enzyme methylation cascade was modified to incorporate radical SAM enzymes. Reactions were performed in an anoxic chamber, and the buffer was complemented with 300 mM NaCl to ensure enzyme stability. Also, 1 mM Ti(III)citrate was added to the assays as a reducing agent as described before.^[56,78]^ Adding cobalamin to the assay mixture was not necessary as the enzyme was reconstituted with hydroxocobalamin after purification, which remains tightly bound to the enzyme.^[56]^ A 24-mer peptide fragment of methyl-coenzyme M reductase was used as the substrate as described previously.^[56]^ Analysis of the linear cascade consisting of MAT, QCMT, and MTAN by liquid chromatography mass spectrometry (LC-MS) revealed full conversion after 20 h (Figure S4a). In the next step, QCMT was coupled to the full SAM regeneration system, initially with a 10× excess of substrate over AMP (0.4 mM and 0.04 mM, respectively), corresponding to 20 possible TTN of the regeneration system as two SAM molecules are required for one methyl transfer to the peptide. After 20 h, analysis by LC-MS showed a conversion of 90% ± 3%, corresponding to a TTN of 18 ± 1 (Figure S4b).

In experiments with 25× excess of peptide over AMP (in theory allowing a TTN of 50), an average conversion of 56% ± 22% was reached, amounting to 28 ± 11 TTN. (Figure 4b) This is less than for MTs and APTs, and might be related to the lower amounts of AMP (0.02 mM vs. 0.04 mM) used due to low concentration of peptide substrate.

### QCMT is able to transfer an ethyl group from SAE via cobalamin on its peptide substrate

Since conventional MTs are able to transfer alternative alkyl chains such as ethyl (from L-Eth/SAE),^[33–35]^ we wondered if radical SAM MTs such as QCMT would be able to generally catalyse alkyl transfer to the peptide substrate. Such an activity by class B radical SAM MTs would be highly interesting, as it would potentially enable the stereo- and regioselective introduction of new functionalities to sp^3^ carbons *via* a radical mechanism.

These modified products would also be useful tools in mechanistic studies on different types of radical SAM enzymes. Nevertheless, the SAM analogue would have to support the generation of the DOA• radical and the corresponding alkylcobalamin. In addition, the alkylcobalamin must be able to react with the substrate radical. The ability to generate the DOA• radical from SAM analogues has been previously demonstrated for *Se*-adenosyl selenomethionine (*Se*SAM), SAE, *S*-adenosyl allylhomocysteine, nucleobase analogues, and the sulfoxide derivative of SAH.^[36–38,79,80]^ Alkylcobalamins other than MeCbl and AdoCbl have been produced by preparative synthetic methods.^[81]^

We investigated the three enzyme cascade consisting of MAT, QCMT and MTAN with L-Eth, D,L-allylhomocysteine and L-propargylselenohomocysteine instead of L-Met, using the *Methanocaldococcus jannaschii* MAT variant L147A/I351A (PC*Mj*MAT) optimised for the formation of SAM-analogues with bulkier residues (Figure 4e).^[82]^ LC-MS analysis of the L-Eth assay showed a signal at 2419.0 Da, corresponding to a mass shift of +28 Da, supporting ethylation of the peptide. In addition, assays containing no MTAN showed peptide ethylation with SAH and DOA formation (Figure 4f, Figure S4c). Further analysis of the cobalamin molecule bound to the enzyme revealed a species with the m/z of 679.8 Da, corresponding to ethylcobalamin + 2 H^+^. Neither allylation, nor propargylation of the peptide was observed. When looking for the corresponding alkylcobalamins, propargylcobalamin + 2 H^+^ could be detected at an m/z of 684.8 Da (Figure 4d), however no allylcobalamin. Analysis of the propargylation assays containing no MTAN revealed a peak likely corresponding to the seleno-analogue of SAH (*Se*SAH), while only trace amounts of DOA could be observed in the chromatogram. Interestingly, even though no allylcobalamin was detected, SAH was observed in the assay with the allyl analogue (Figure 4f; Figure S4c). The lack of DOA formation for both, the allyl and propargyl analogue, suggests that either these analogues are not able to support radical generation due to steric restrains, or that the *S/Se*-alkyl bond is cleaved rather than the *S/Se*-C5′ bond. The latter could be supported by the π-system of these analogues, as it makes the *S/Se*-alkyl bond the weakest bond. However, experiments with allyl-SAM and NosN resulted in functional enzyme with cleavage of the S-C5′ bond.^[37]^ In addition, photoinduced cleavage of SAM bound to HydG or PFL-AE suggests that regioselectivity of bond cleavage is determined by the geometry of SAM bound to the [4Fe4S]-cluster.^[83]^ Hence, the bulkier residues would have to alter the binding of SAM to allow cleavage of the S/Se-alkyl bond. Interestingly, the expected C-alkyl bond cleavage byproduct (Se)SAH is formed with all tested analogues. Nevertheless, it is also the byproduct of cobalamin alkylation and therefore an expected byproduct for the enzyme. The lack of allylcobalamin can be explained with rapid degradation by homolytic cleavage of the allyl-Co bond under formation of a stable allyl radical and cob(II)alamin (Figure S4e).

To investigate if the *Se*-*C*-bond hinders the formation of the DOA• radical in the propargylation assays, L-selenomethionine was used to yield *Se*SAM. Methylation of the peptide indicated a fully functional enzyme. As HPLC analysis confirmed the formation of DOA (Figure S4c), we concluded that the *Se*-*C*-bond was not impeding the radical formation.

Finally, we investigated whether the SAM regeneration system was able to support ethylation of the peptide substrate. L-Met was replaced with L-Eth and the peptide substrate was present in 5× excess. After 20 h, a portion of approximately 7% ethylated product was detected (Figure 4c), which was in the same range as the conversion obtained for the linear cascade (∼14%, Figure S4d).

The incorporation of radical SAM enzymes is promising for future applications due to their vast chemical capabilities. Particularly in drug synthesis the ever expanding chemical space of radical SAM enzymes can be useful for challenging reactions, e. g. stereo- and regioselective methylation of sp^3^ carbons. In addition to the possibility of enhancing the diversity of radical SAM MT products by transferring alternative alkyl chains, this might be interesting in terms of using SAM analogues for mechanistic studies as has been shown before.^[36,37,79,80]^ Together with the increasing number of solved enzyme structures, it might be possible to design enzyme variants with a broader or altered substrate range, as has been shown for MATs and conventional MTs.^[82,84–86]^

### SAM Regeneration from MTA Supports Aminopropyl Transfer in Polyamine Synthesis

Finally, the flexibility of the system was investigated by integrating polyamine synthesis. The transfer of the aminopropyl unit in the biosynthesis of polyamines from SAM is a two-step process: first, SAM is decarboxylated by SAM-decarboxylase (SAMDc, E.C. 4.1.1.50). Then, the generated decarboxy-SAM (dcSAM, *S*-adenosylmethioninamine) is used as an aminopropyl group donor, with e.g. putrescine attacking dcSAM in an S_N_2 reaction as a nucleophile. This polyamine aminopropyltransferase (PAPT) catalysed transfer produces MTA as a byproduct (Figure 1a).^[87]^ Hence, SAM regeneration from MTA is required (Figure 5).

**Figure 5.**
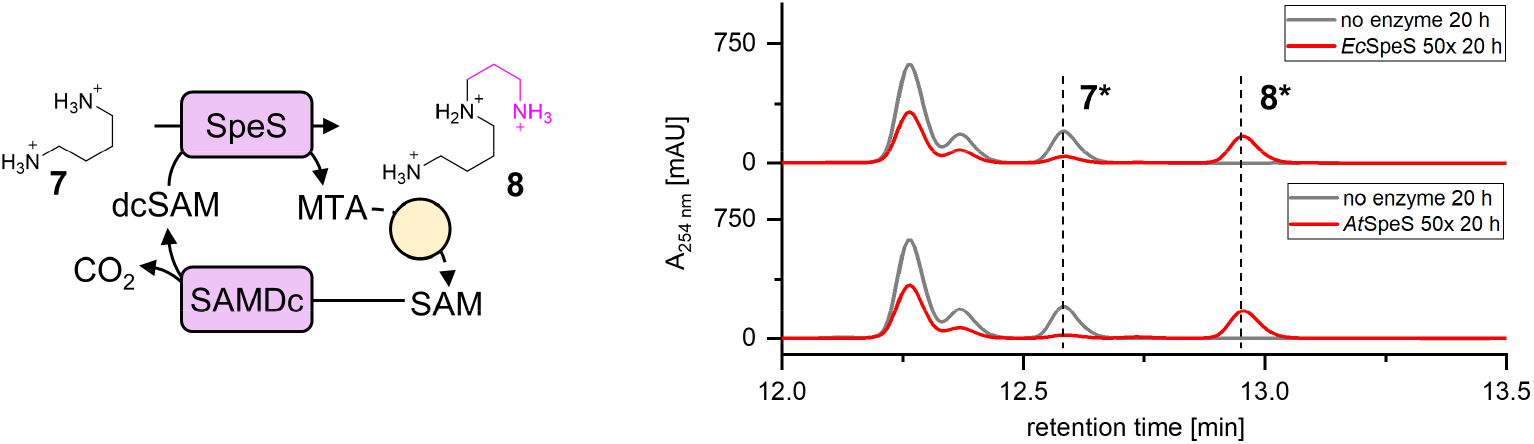
The SAM regeneration system with APTs. Spermidine synthase (SpeS) alongside SAM decarboxylase (SAMDc) was utilised as a model enzyme for aminopropyl transfer from dcSAM to putrescine (**7**) yielding spermidine (**8**). The SAM regeneration system is indicated by the yellow circle. The top chromatogram shows conversion for *Ec*SpeS, the bottom chromatogram shows conversion for *At*SpeS, both at 50× excess of substrate over AMP. All peaks correspond to the BzCl-derivatised polyamines (**\***).

As PAPT model enzymes, spermidine synthase (SpeS, E.C. 2.5.1.16) from both *Arabidopsis thaliana* (*At*SpeS isoenzyme 1)^[88,89]^ and *E. coli* (*Ec*SpeS)^[87]^ were selected using putrescine as a substrate. SAMDc from *Thermotoga maritima* (*Tm*SAMDc) was used for the first step of the reaction.^[90]^ For HPLC analysis with detection at 254 nm, the assay samples were derivatised with benzoyl chloride (BzCl).^[91]^

The envisioned multi-enzyme system was first investigated using a linear SAM supply cascade consisting of *Ec*MAT, *Tm*SAMDc, *At*SpeS (or *Ec*SpeS), and *Ec*MTAN. To avoid derivatisation of Tris, the buffer system was altered to MOPS for all assays containing polyamines (Figure S5a, Figure S5b). As expected,^[92]^ the removal of inhibitory MTA by addition of MTAN to the experiment accelerated the formation of spermidine. For *Ec*SpeS, addition of MTAN increased the conversion after 1 h by 25%, for *At*SpeS by 16% (Figure S5c). Interestingly, further aminopropylation of spermidine yielding spermine was detected only in assays with MTAN present after 20 h, supporting the inhibitory character of MTA on SAMDc and/or SpeS. Formation of spermine was observed for both *At*SpeS and *Ec*SpeS (Figure S4d); while this has been known for *Ec*SpeS, *At*SpeS has previously been reported to lack spermine synthase activity in *in vivo* experiments.^[88]^ In the next step, the ATP production system was added to close the cycle. The performance of the system with SAMDc and SpeS was evaluated after 20 h. Although the system gets more complex through the introduction of another enzyme (SAMDc), the performance was comparable to conventional MTs. Similar conversion rates for both SpeSs were observed, with >94% conversion (putrescine to spermidine and spermine) for a 10× excess of substrate over AMP and conversion of up to 57% for 100× excess of substrate (Table 1, Figure 5, Figure S5d). Conversion to spermine was only observed for samples with 0.2 mM AMP, suggesting that a high level of spermidine must be available as substrate for spermine synthesis (Figure S5d).

## Conclusion

*In situ* synthesis and regeneration of SAM or SAM analogues are important steps towards the efficient utilisation of the cofactor for biocatalytic applications. All SAM regeneration systems designed so far have been limited to SAH producing MTs.^[61–63]^ The SAM regeneration system presented here unlocks the full potential of the highly versatile cofactor SAM, including radical SAM enzymes and polyamine synthases. This is achieved by regenerating SAM *via* the former dead-end byproduct adenine using stable starting materials and catalytic amounts of AMP. As SAH, MTA and DOA have been shown to be potential inhibitors for SAM-dependent enzymes^[41,57,92]^, their removal and regeneration represent a valuable tool to increase the activity of SAM-dependent enzymes for synthetic and mechanistic studies. Balancing the system regarding enzyme concentrations might further increase conversion at low AMP levels, as there are three enzymes competing for ATP present (RK, RPPK and MAT); nevertheless, the minimal amount of cofactor building block will be dictated by the enzymes’ intrinsic kinetic parameters (Table S7). Furthermore, the accumulation of the ribosyl byproduct of the MTAN reaction could be prevented by further degradation. While none of the three byproducts are known inhibitors or substrate of the RK or RPPK, it is still possible that these compounds influence the system.

The supply and regeneration systems are able to support *in situ* synthesis of alkyl analogues and alkyl transfer as established for conventional MTs and shown here for the radical SAM QCMT. This opens up the possibility of directly introducing desired functional groups stereo- and regioselectively to sp^3^ carbon atoms enzymatically *via* a radical mechanism. Building up the SAM molecule from the basic building blocks nucleobase, sugar, phosphates, and alkylated homocysteine gives full flexibility for all types of SAM analogues.^[34,93]^

## Acknowledgements

We thank Katharina Strack (University of Freiburg) for her skilful technical assistance. Jaspreet Chahal (University of Southampton) and Sonya Danesh (University of Freiburg) are acknowledged for help with optimisation of *rppk* production, and Dr Manfred Keller and Dr Thomas Haas (University of Freiburg) for help with NMR measurements. We also thank Dr Peter Roach for helpful advice on radical SAM chemistry. Dr Florian Hubrich (ETH Zurich) is thanked for proofreading and helpful discussions.

This work was funded by the European Research Council (ERC project 716966; L.G., D.M., L.K., J.N.A.), Deutsche Forschungsgemeinschaft (RTG1976; L.K., J.G., G.L., J.N.A.; IRTG2027 N.C., A.R.). X.W. is the recipient of a PhD stipend from the Camille Henry Dreyfus Foundation.

