## Supplemental Material for "Biomimetic S-Adenosylmethionine Regeneration Starting from Different Byproducts Enables Biocatalytic Alkylation with Radical SAM Enzymes"

[e] Dr. S. Mordhorst
Pharmaceutical Institute, Pharmaceutical Biology
University of Tübingen
Auf der Morgenstelle 8, 72076 Tübingen (Germany)

[f] Prof. Dr. H. J. Jessen
Institute of Organic Chemistry
University of Freiburg
Albertstr. 21, 79104 Freiburg (Germany)

### Experimental Procedures (and Tables S1 to S6)

### Materials & Reagents

All chemicals were purchased in the highest purity available from Sigma-Aldrich (ATP, ADP, AMP, SAM, SAH, MTA, l-methionine, l-ethionine, l-selenomethionine, *N*-ethylanthranilic acid, 3,4-dihydroxybenzoic acid, putrescine, spermidine, spermine), Acros Organics (isovanillic acid and sodium polyphosphate), Alfa Aesar (*N*-methylanthranilic acid, vanillic acid), AppliChem (adenine) and Fluka (anthranilic acid). Ingredients for buffers and cultivation media were purchased from Carl Roth. The peptide substrate for QCMT (*H*-PTMMEDHFGGSQRAGVIAAASGLS-*OH*) was purchased at TAG Copenhagen (Denmark).

**Table S1** UniProt accession numbers and calculated extinction coefficients (ExPASy ProtParam)[1] of enzymes used in this study.

| Enzyme | UniProt accession number | $\varepsilon_{280}$ [L·mol$^{-1}$·cm$^{-1}$·1000$^{-1}$] |
|---|---|---|
| *Ec*RK | P0A9J6 (RBSK_ECOLI) | 12.49 |
| *Mt*RPPK | P9WKE3 (KPRS_MYCTU) | - |
| *Ec*RPPK | P0A717 (KPRS_ECOLI) | 6.21 |
| *Ec*APRT | P69503 (APT_ECOLI) | 10.93 |
| *Sm*PPK2 | Q92SA6 (Q92SA6_RHIME) | 57.41 |
| *Aj*PPK2 | Q83XD3 (Q83XD3_ACIJO) | 121.48 |
| *Ec*MAT | P0A817 (METK_ECOLI) | 40.13 |
| *Tk*MAT | Q5JF22 (METK_THEKO) | 30.83 |
| *Mj*MAT* | Q58605 (METK_METJA) | 25.12 |
| *Ec*MTAN | P0AF12 (MTNN_ECOLI) | 6.21 |
| *Rn*COMT | P22734 (COMT_RAT) | 30.62 |
| *Rg*ANMT | A9X7L0 (ANMT_RUTGR) | 31.78 |
| *Tm*SAMDc | Q9WZC3 (SPEH_THEMA) | 20.07 |
| *Ec*SpeS | P09158 (SPEE_ECOLI) | 32.24 |
| *At*SpeS | Q9ZUB3 (SPDS1_ARATH) | 35.41 |
| QCMT | A0A1G8Z8Q2 (A0A1G8Z8Q2_9EURY) | - |

**Mj*MAT L147A/I351A was used.

### Methods

#### Protein Production

The genes encoding ribokinase (RK) and adenosine phosphoribosyltransferase (APRT) were amplified from *E. coli* genomic DNA by PCR using the primers given in Tab. 2 and cloned into pET28a using T4 DNA ligase (New England Biolabs GmbH, Germany). The gene encoding ribose-phosphate pyrophosphokinase (RPPK) was also amplified from genomic DNA of *E. coli* and cloned into pET28a using In-Fusion cloning (Takara Bio Europe, France).

**Table S2** PCR primers used in this study. Restriction sites are shown in bold.

| Primer | Sequence (5' → 3') |
|---|---|
| *ecrk_for* | GCGGCAGC**CATATG**ATGCAAAACGCAGGCAGCC |
| *ecrk_rev* | GCGGCCGC**AAGCTT**TCACCTCTGCCTGTCTAAAAATGCG |
| *ecaprt_for* | GCGGCAGC**CATATG**ATGACCGCGACTGCACAGC |
| *ecaprt_rev* | TGGTGGTG**CTCGAG**TTAATGGCCCGGGAACGG |
| *ecrppk_for* | CGCGCGGCAGC**CATATG**CCGGATATGAACC |
| *ecrppk_rev* | GTGCGGCCGC**AAGCTT**AATGTTCAAACATC |

QCMT was produced and purified as described previously.[2] All other enzymes were produced as follows: *E. coli* BL21-Gold(DE3) (Agilent, Santa Clara, USA) cells transformed with the respective pET28a construct were grown in lysogeny broth (LB) medium supplemented with kanamycin (50 µg/mL) at 37 °C with shaking until an $OD_{600}$ of 0.5-0.6 was reached. Gene expression was induced by addition of 0.2 mM of isopropyl β-D-1-thiogalactopyranoside (IPTG) and cultures were further incubated overnight at 20 °C. The harvested cell pellet was resuspended in lysis buffer [40 mM Tris-HCl, pH 8.0, 100 mM NaCl, 10% (*w*/*v*) glycerol] and lysed by sonication (Branson Sonifier 250, Emerson, USA) The lysates were used for protein purification by Ni(II)-NTA affinity chromatography (Invitrogen, Thermo Fisher Scientific, USA) as previously described.[3] Desalting was carried out using a PD-10 column (GE Healthcare Life Sciences, UK). For PC*Mj*MAT the protein was further purified by heating at 80 °C for 20 min followed by 30 min centrifugation at 18213 × g (Eppendorf centrifuge 5427 R). Protein concentrations were determined by UV/vis absorption with calculated extinction coefficients ($\varepsilon_{280}$, ExPASy ProtParam)[1] using a NanoDrop 2000 (Thermo Fisher Scientific, USA). Concentrations of *Mt*RPPK and QCMT were determined by Bradford analysis.[4] Purity of the enzymes was assessed by SDS-PAGE (Fig. S1).[5]

**Synthesis of methionine analogues**

Allyl-D,L-homocysteine

As described previously,[6] D,L-homocysteine (100 mg, 0.74 mmol, 1 eq.) was dissolved in 2 mL of $H_2O$ and 124 mg of $NaHCO_3$ (1.48 mmol, 2 eq.) were added. To this solution 115 µL of allyl bromide (1.34 mmol, 1.8 eq.) was added and the mixture was stirred for 3 h at 40 °C. The reaction was cooled to room temperature, adjusted to pH = 3 with 1 M HCl and purified via preparative HPLC. The product was obtained as white crystalline solid (56 mg, 0.32 mmol, 43 %). The identity was confirmed by HRMS (LTQ Orbitrap XL, Thermo Fisher Scientific, Fig. S6), $^1$H NMR and $^{13}$C NMR (Fig. S7).

**$^1$H NMR** (400 MHz, $D_2O$ + 1 % NaOD) δ (ppm) = 5.82 (m, 1H), 5.31 – 5.10 (m, 2H), 3.30 (m, 1H), 3.20 (m, 2H), 2.53 (m, 2H), 1.98 – 1.67 (m, 2H).

**$^{13}$C-{$^1$H}-NMR** (101 MHz, $D_2O$ + 1 % NaOD) δ (ppm) = 182.8, 134.0, 117.6, 55.3, 34.4, 33.5, 26.3.

Propargyl-L-selenohomocysteine

First, L-selenohomocysteine was synthesized in two steps starting from L-homoserine (Activate Scientific) as previously reported by Bothwell *et al.*.[7] Under an argon atmosphere L-selenohomocysteine (119 mg, 0.65 mmol, 1 eq.) was dissolved in 20 mL dry ethanol. Sodium borohydride (124 mg, 3.28 mmol, 5 eq.) was added and the mixture was stirred for 15 min at room temperature. Sodium bicarbonate (150 mg, 1.78 mmol) and propargyl bromide (80 % in toluene, 156 µL, 208 mg, 1.26 mmol, 2 eq.) were added and the mixture stirred for 16 h. The solvent was removed under reduced pressure. The raw product was dissolved in 240 mM HCl (10 mL) and purified *via* preparative HPLC. The product was obtained as white crystalline solid (100mg, 0.45 mmol, 69 %). The identity was confirmed by HRMS (LTQ Orbitrap XL, Thermo Fisher Scientific, Fig. S8), $^1$H NMR and $^{13}$C NMR (Fig. S9).

**$^1$H NMR** (400 MHz, $D_2O$) δ (ppm) = 3.84 (t, $J$ = 6.9, 5.6 Hz, 1H), 3.39 – 3.28 (m, 2H), 2.92 – 2.82 (m, 2H), 2.67 (t, $J$ = 2.7 Hz, 1H), 2.38 – 2.19 (m, 2H).

**$^{13}$C-{$^1$H}-NMR** (101 MHz, $D_2O$) δ (ppm) = 171.8, 79.2, 69.9, 52.5, 28.9, 16.7, 4.2.

**Enzyme assays**

In general, all enzyme assays were at least performed in triplicates. If not stated differently, negative controls contained no enzyme.

Ribose phosphorylation

Initial assays investigating the phosphorylation of ribose towards PRPP were set up with *Ec*RK, *Mycobacterium tuberculosis* RPPK (*Mt*RPPK)[8], *Sm*PPK2 and *Aj*PPK2 with an excess of polyphosphate (polyP) and catalytic amounts of AMP. The assay composition was as follows:

50 mM Tris buffer pH 7.5, 50 mM KCl, 20 mM $MgCl_2$, 30 mM polyP, 3 mM D-ribose, 0.03 µM *Ec*RK, 2 µM *Mt*RPPK, 1 µM *Aj*PPK2 and 1 µM *Sm*PPK2. $D_2O$ was added to a final concentration of 10% and 0.5 mM cAMP was added as an internal standard. The assays were started by addition of 0.2 mM AMP. PRPP production was followed by $^{31}P$ NMR on a Bruker (Billerica, USA) Avance Neo 400 MHz spectrometer at 162 Mhz with a 5 mm BB Prodigy probehead. The sample was tempered to 37 °C without agitation. The spectrum of the last data point was manually phased, the resulting parameters were applied to all other datasets. PRPP concentration was determined by integration of the anomeric-bound pyrophosphate unit with signals at -11.5 ppm (α-phosphate), at -5.80 to -5.86 ppm (terminal β-phosphate) and at 3.27 to 2.87 ppm (5'-phosphate). D-Ribose-5-phosphate concentration was determined by integration of signals shifted from 3.28 ppm to 2.87 ppm.
Assays investigating the nucleotide production from D-ribose were carried out as follows: 50 mM $KP_i$ buffer (pH 7.5), 50 mM KCl, 20 mM $MgCl_2$, 1 mM adenine, 1 mM D-ribose, 20 mM polyP, 2.9 µM *Ec*RK, 2.7 µM *Mt*RPPK, 4.5 µM *Ec*APRT, 0.9 µM *Aj*PPK2, 1.4 µM *Sm*PPK2 and 0.2 mM ATP were added to a final volume of 1 mL and incubated for 1 h at 37 °C without agitation. Samples were taken at 0 min, 15 min, 30 min, and 60 min and snap frozen in liquid nitrogen. For HPLC analysis the samples were centrifuged for 20 min at 4 °C, 18213 × g (Eppendorf centrifuge 5427 R) and the supernatant was filtered using Vivaspin® 500 Centrifugal Concentrators (10,000 MWCO, Sartorius AG, Germany). Samples were analysed using HPLC method A.

Methyl Transferase Assays

The assays for SAM regeneration were carried out in a reaction mixture (400 µl) with final concentrations of 50 mM Tris buffer (pH 7.5) with 20 mM $MgCl_2$, 50 mM KCl and 20 mM polyP. The substrates for the methylation reactions, either anthranilic acid or 3,4-dihydroxybenzoic acid, were added to the final concentration of 2 mM. D-ribose and L-methionine (L-Met) were added in slight excess at 3 mM. For ethylation assays, L-ethionine (L-Eth) was used instead of L-Met (also 3 mM). The enzymes were used at the following final concentrations: 3 µM *Ec*RK, 3 µM *Ec*RPPK, 3 µM *Ec*APRT, 1 µM *Aj*PPK2, 1 µM *Sm*PPK2, 10 µM *Ec*MAT and 1 µM of *Ec*MTAN. Either 10 µM *Rn*COMT or 10 µM *Rg*ANMT. For ethylation assays, *Ec*MAT was replaced with *Tk*MAT. Varying final concentrations (0.2 mM, 0.04 mM and 0.02 mM) of the cofactor building block AMP were added to achieve maximum possible turnover numbers of 10, 50 and 100, respectively. The reaction mixtures were incubated at 37 °C, 350 rpm for 20 h. The reaction was quenched by the addition of $HClO_4$ to a final concentration of 2% and samples were snap frozen in liquid nitrogen. Samples for HPLC analysis were prepared as described above and analysed by HPLC method B.

Spermidine Synthase Assays

Prior to all assays, *Tm*SAMDc was activated by heating at 80 °C for 1 h.[9] Initial polyamine synthesis assays were set up in the linear SAM supply cascade containing MAT and MTAN. 50 mM MOPS, Tris or HEPES buffer pH 7.5, 50 mM KCl, 20 mM $MgCl_2$, 3 mM L-Met, 3 mM ATP, 2 mM putrescine, 10 µM *Ec*MAT, 1.5 µM *Ec*MTAN, 20 µM SpeS and 40 µM *Tm*SAMDc were added to a final volume of 500 µL. The assays were incubated for 20 h at 37 °C.

Polyamine synthesis in the SAM regeneration system was set up as described above for ANMT and COMT with the following alterations: The buffer was 50 mM MOPS pH 7.5 and the MT was replaced by 40 µM *Tm*SAMDc and 20 µM SpeS (*Ec*SpeS or *At*SpeS).

For all SpeS assays, samples were immediately snap frozen in liquid nitrogen without prior quenching and stored at -20 °C until further preparation. The frozen samples were centrifuged at 4 °C at 18213 × g until thawed. 20 µL of the supernatant was used for sample preparation. 20 µL NaOH (0.2 M) and 20 µL benzoyl chloride (40 mM) were added to the sample prior to incubation at 25 °C, 350 rpm for 30 min. The derivatisation reaction was quenched by the addition of 60 µL formic acid (2.5%). Standards for qualitative and quantitative analysis were derivatised accordingly. The derivatised samples were prepared for HPLC analysis (Method C) by centrifugation at 4 °C, 18213 × g for 45 min.

Class B radical SAM MT Assays

All assays containing QCMT were conducted under strict exclusion of oxygen in a Coy anaerobic chamber maintained at 97.5% $N_2$ and 2.5% $H_2$. Except for the enzymatic preparations, peptide and Ti(III)citrate, the stock solutions for all other reaction components were prepared freshly inside the anaerobic chamber.

Assays using the linear SAM supply cascade contained 50 mM Tris buffer (pH 7.5) supplemented with 300 mM NaCl, 20 mM $MgCl_2$ and 50 mM KCl. In addition, 1 mM Ti(III)citrate, 0.05 mM peptide (24mer, *H*-PTMMEDHFGGSQRAGVIAAASGLS-*OH*), 2 mM L-Met and 2 mM ATP were added. For the methylation cascade 10 µM *Ec*MAT, 10 µM QCMT and 1 µM *Ec*MAT were used. For assays containing methionine analogues, L-Met was replaced with either 3 mM L-selenomethionine, 3 mM L-Eth, 6 mM D/L-allylhomocysteine or 3 mM L-propargyl-selenohomocysteine and 50 µM PC*Mj*MAT replaced *Ec*MAT.[10] Assays without *Ec*MTAN were conducted for the analysis of SAH and DOA. All assays were incubated for 20 h at 350 rpm, 37 °C. Samples for peptide, SAH and DOA analysis were quenched by addition of $HClO_4$ at a final concentration of 2.5%. The vials were immediately snap-frozen and stored at -20 °C until use. For LC-MS analysis, the samples were thawed on ice and centrifuged (18213 × *g*, 20 min, 4 °C). The supernatant was passed through a syringe filter (0.2 µm cut-off) and the flow-through obtained was used for LC-MS analysis. Samples for the analysis of cobalamin bound to the enzyme were prepared as previously described.[2]

Assays for investigating the SAM regeneration system with QCMT were carried out in 50 mM Tris buffer (pH 7.5) containing 300 mM NaCl, 20 mM $MgCl_2$ and 50 mM KCl. To this, L-Met, D-ribose, and polyP were added in excess at 3 mM, 3 mM and 20 mM, respectively. Peptide (24mer) was added at 0.5 mM and Ti(III)citrate at 2 mM of final concentration. *Ec*RK, *Ec*RPPK, *Ec*APRT, *Sm*PPK2, *Aj*PPK2, *Ec*MAT and *Ec*MTAN were added as described above. The reaction mixtures were incubated at 37 °C, 350 rpm for 20 h. Assays containing L-Met analogues were altered as described above for the linear cascade. The samples were prepared for LC-MS analysis as described above.

**HPLC analysis**

Method A: Analysis of samples from assays investigating the formation of nucleotides from D-ribose and adenine was conducted by a reversed phase HPLC method adapted from LIU *et al.*[11] An Agilent Technologies 1260 infinity II HPLC system with an ISAspher 100-5 C18 column (250 mm × 4.0 mm, 5 µm, ISERA GmbH, Germany) was used. The temperature of the column was held at 25 °C. The elution gradient was a mixture of two solvents: 100 mM $KP_i$ buffer at pH 7.0 (A) and $CH_3CN$ (B). Following gradient was applied: 0-3 min 0% B, 3-5 min 0% to 5% B, 5-7 min 5% to 20% B, 7-8.3 min 20% to 25% B, 8.3-9 min 25% to 0% B and 9-15 min 0% B. The sample injection volume was 10 µL and the flow rate throughout the 15 min run was 1.2 mL/min. Detection of analytes was performed at 254 nm. Retention times of standards are given in Tab. 3.

**Table S3** Retention times of standards determined for HPLC method A.

| **Substance** | **Retention time [min.]** |
|---|---|
| Adenine | 9.0 |
| ADP | 7.7 |
| AMP | 8.0 |
| ATP | 6.7 |

Method B: Analysis of substrates and products of methyltransferase reactions was carried out on an Agilent Technologies 1260 infinity II HPLC system using ISAspher 100-5 C18 column (125 mm × 4.0 mm, 3 µm, ISERA GmbH, Germany) using a mobile phase of (A) 0.1% formic acid in $H_2O$ and (B) 100% $CH_3CN$ at a flow rate of 1 mL/min. The column was equilibrated with 3% solvent B and after injection of 10 µL of sample, solvent B was applied with a following gradient: 0-4 min 3% B, 4-7 min 3% to 70% B, 7-7.10 min 70% to 100% B, 7.10-10 min 100% B, 10-11 min 100% to 3% B, 11-14 min 3% B. Detection of substrates and methylated products was performed at 254 nm with a diode array detector (DAD). Quantitative determination of substrates and products was carried out by recording calibration curves using authentic references of the following substrates and methylated products: anthranilic acid, 3,4-dihydroxybenzoic acid, *N*-methylanthranilic acid, *N*-ethylanthranilic acid, vanillic acid and isovanillic acid. The retention times and calibration curve equations are given in Tab. 4.

**Table S4** Retention times of standards determined for HPLC method B.

| **Substance** | **Retention time [min.]** | **Calibration curve** | **$R^2$** |
|---|---|---|---|
| 3-4-dihydroxybenzoic acid | 7.1 | y = 2032.9x | 0.99 |
| Vanillic acid, isovanillic acid | 7.6 | y = 2388.1x | 0.95 |
| Anthranilic acid | 8.0 | y = 1250.1x | 0.99 |
| *N*-Methylanthranilic acid | 8.7 | y = 1990.8x | 0.98 |
| *N*-Ethylanthranilic acid | 8.9 | y = 2151.6x | 0.99 |

Method C: Derivatised polyamine samples were analysed by reversed phase HPLC on an Agilent 1100 Series with an ISAsphere 100-5-C18 column (250 mm × 4.0 mm, 5 µm, ISERA GmbH, Germany). Mobile Phase A was water with 1% $CH_3CN$ and 0.1% formic acid, mobile phase B was $CH_3CN$. The following gradient was applied at a flow rate of 1.0 mL/min: 0-1.5 min 2% B, 1.5-8 min 2% to 33% B, 8-11 min 33% to 95% B, 11-15.5 min 95% B, 15.5-17.5 min 95% to 2% B, 17.5-22 min 2% B. Injection volume was 10 µL. Peaks were detected at 254 nm by DAD. The retention times an calibration curves are given in Tab. 5.

**Table S5** Retention times of standards determined for HPLC method C.

| **Substance** | **Retention time [min.]** | **Calibration curve** | **$R^2$** |
|---|---|---|---|
| Benzoic acid | 9.0 | n. d. | n. d. |
| Putrescine* | 12.6 | y = 450.44x | >0.99 |
| Spermidine* | 13.0 | y = 529.18x | >0.99 |
| Spermine* | 13.2 | y = 599.99x | >0.99 |
| Tris* | 12.8 | n. d. | n. d. |

Method D: Analysis of QCMT reaction byproducts was conducted by reversed phase HPLC on an Agilent Technologies 1260 infinity II HPLC system with an ISAspher 100-5 C18 column (250 mm × 4.0 mm, 5 µm, ISERA GmbH, Germany). Mobile phase A was 40 mM Na acetatae buffer pH 4.2, mobile phase B was $CH_3CN$. The gradient was set up as follows at a flow rate of 0.5 mL/min: 0-16 min 2% B to 30% B, 16-18 min 30% B, 18-20 min 30% to 2% B, 20-30 min

2% B. Injection volume was 10 µL, peaks were detected using a DAD at 260 nm. Retention times of standards are given in Tab. 6.

**Table S6** Retention times of standards determined for HPLC method D.

| **Substance** | **Retention time [min.]** |
|---|---|
| Adenine | 13.1 |
| ADP | 9.0 |
| AMP | 11.4 |
| ATP | 6.2 |
| DOA | 16.6 |
| MTA | 19.4 |
| SAH | 13.7 |
| SAM | 10.8 |

**LC-MS Analysis**

Peptide Analysis

Analysis of the 24mer peptide was conducted as previously described.[2] Quantitative peptide turnover measurements were performed with the UniDec Software using the interpolate function.[12] Masses were calculated and measured as average isotopic masses when not stated different.

Cobalamin analysis

Analysis by LC-MS of cobalamin bound to QCMT was conducted as previously described.[2] Masses were calculated and measured as average isotopic masses when not stated differently.

### Supplementary Data

**Table S7** Kinetic parameters of enzymes used in this study. As no parameters were found in literature for *Sm*PPK2 and *Aj*PPK2, PPK2 from *Francisella tularensis* (*Ft*PPK2) and *Paenarthrobacter aurescens* (*Pa*PPK2) are listed for coparison.

| Enzyme | Substrate | $K_M$ [mM] | Conditions | Reference |
|---|---|---|---|---|
| *Ec*RK | D-Ribose | 0.18 | pH 7.4, 37 °C | [13] |
| *Ec*RPPK | PRPP | 0.19 | pH 8.0, 37 °C | [14] |
| | ATP | 0.023 | pH 8.0, 37 °C | [15] |
| *Ec*APRT | Ade | 0.02 | not specified | [16] |
| | PRPP | 0.125 | 2x excess $Mg^{2+}$ | [16] |
| *Sm*PPK2 | - | - | - | - |
| *Ft*PPK2 | ADP | 0.546 | pH 8.0, 37 °C | [17] |
| *Aj*PPK2 | - | - | - | - |
| *Pa*PPK2 | AMP | 0.83 | pH 8.0, 30 °C | [18] |
| *Ec*MAT | ATP | 0.17 | pH 8.0, 37 °C | [19] |
| *Ec*MTAN | SAH | 1.3 | pH 7.5, 25 °C | [20] |
| | MTA | 0.43 | pH 7.0, 37 °C | [20] |

### SDS-PAGE analysis

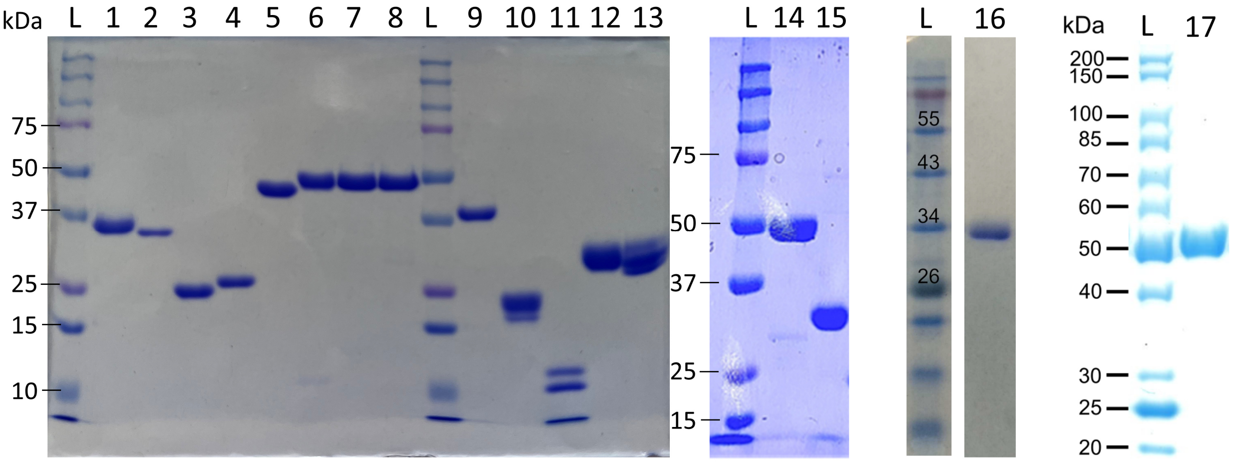

**Figure S1** SDS-PAGE analysis of produced enzymes. 2 µM each were applied. L – protein ladder (Precision Plus Protein Dual Color Standards, Bio-Rad, U.S.A.), 1 – *Ec*RK (32 kDa), 2 – *Ec*RPPK (34 kDa), 3 – *Ec*APRT (22 kDa), 4 – *Ec*MTAN (27 kDa), 5 – *Ec*MAT (44 kDa), 6 – *Tk*MAT (47 kDa), 7 – PC*Mj*MAT (47 kDa), 8 – PC*Mj*MAT (heated, 47 kDa), 9 – *Rg*ANMT (42 kDa), 10 – *Rn*COMT (27 kDa), 11 – *Tm*SAMDc (16 kDa), 12 – *Ec*SpeS (35 kDa), 13 – *At*SpeS (35 kDa), 14 – *Aj*PPK2 (58 kDa), 15 – *Sm*PPK2 (37 kDa), 16 – *Mt*RPPK (38 kDa), 17 – QCMT (49 kDa). The $His_6$-tag is included in all molecular weight calculations.

***In situ* production of PRPP enables the production of ATP from adenine.**

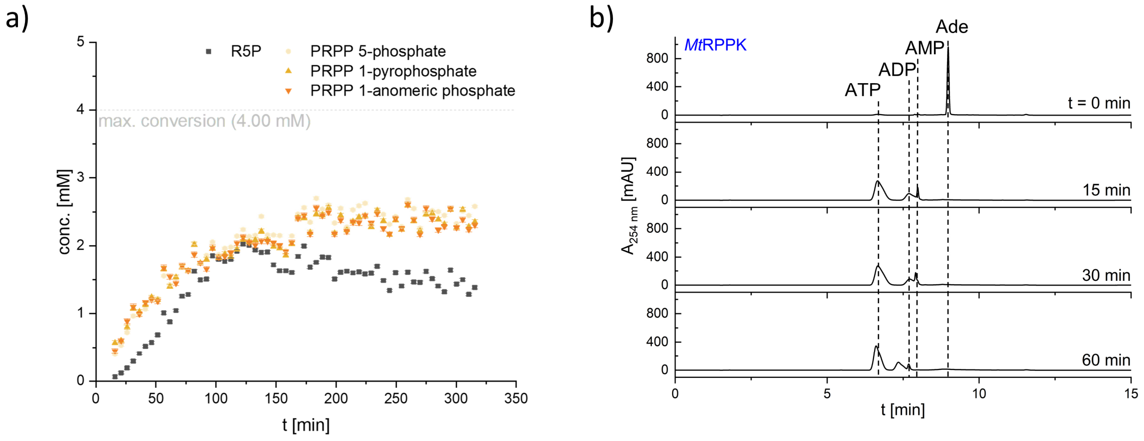

**Figure S2 a)** ATP regeneration for *Ec*RK and *Mt*RPPK using *Aj*PPK2 and *Sm*PPK2 enzymes. PRPP production from D-ribose (3 mM) and a catalytic amount of AMP (0.2 mM) was set up to ensure that the enzymes are active under the conditions established for SAM supply cascade enzymes. The reaction was fuelled by an excess of PolyP. PRPP (orange) and R5P (grey) formation over time was visualised by $^{31}$P NMR. **b)** Adenine (Ade) salvage *via* the ribose phosphorylation cascade containing *Ec*RK, *Mt*RPPK, *Ec*APRT, *Sm*PPK2 and *Aj*PPK2 after t = 0 min, 15 min, 30 min and 60 min (from top to bottom). In addition to D-ribose and Ade (1 mM each), a catalytic amount of ATP (0.2 mM) was used for the RK and RPPK reaction, as well as an excess of polyP for ATP regeneration (Figure 2, dark grey box). Analysis by HPLC at 254 nm showed that a mixture of the nucleotides AMP, ADP and ATP was formed, while no Ade was detected after 15 min. This is consistent with the expected formation of AMP *via* PRPP catalysed by APRT. The distribution between AMP, ADP and ATP results from the activities of the PPK2 enzymes, which are expected to produce a mixture of nucleotides when both enzymes are present, also due to their adenylate kinase reactivity with the resulting formation of ATP and AMP from ADP and *vice versa*.[21]

### SAM regeneration from SAH *via* adenine/MTAN is functional with conventional MTs.

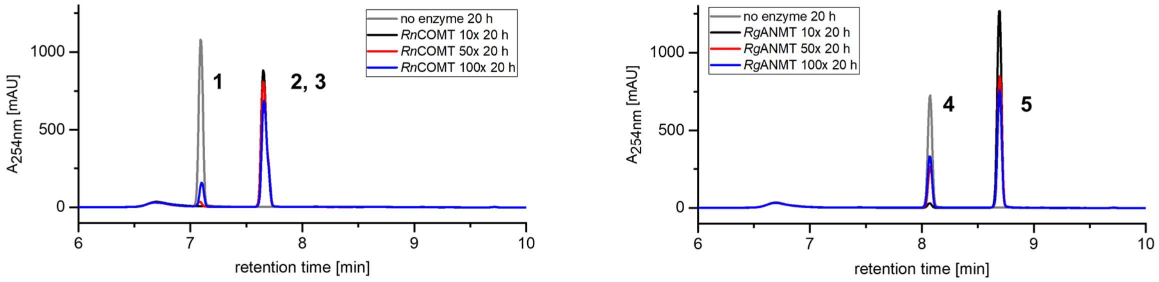

**Figure S3** Chromatograms of typical SAM regeneration assays containing either a substrate excess of 10× (black), 50× (red) or 100× (blue) over AMP. All samples were taken after 20 h. Assays with *Rn*COMT (left) showed conversion of DHBA (**1**) to a mixture of isovanillic acid (**2**) and vanillic acid (**3**) compared to a no enzyme control (grey). Assays with *Rg*ANMT (right) showed conversion of anthranilic acid (**4**) to *N*-methylanthranilic acid (**5**).

### The SAM regeneration system supports Class B radical SAM MTs by SAM regeneration from SAH and DOA.

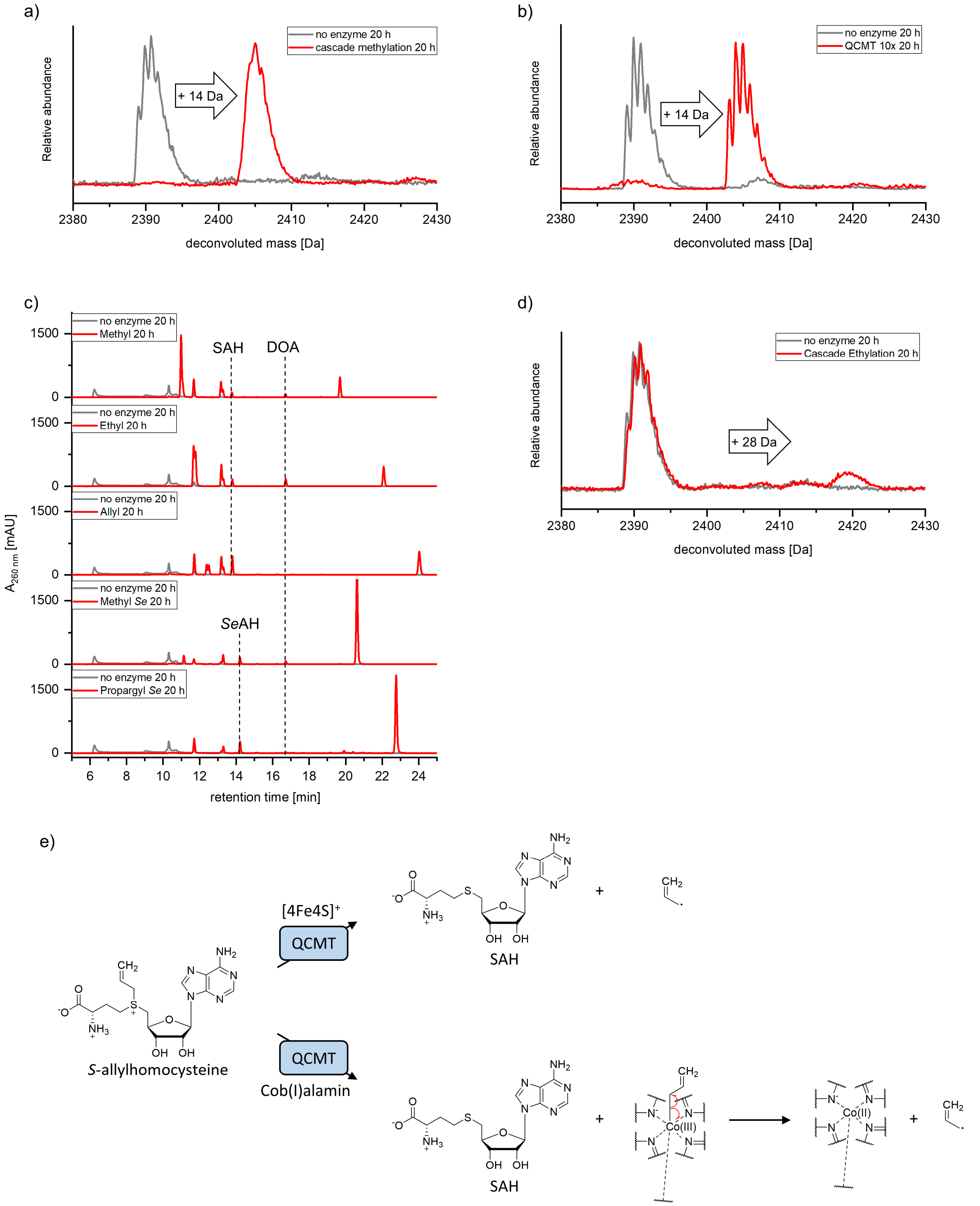

**Figure S4 a)** LC-MS analysis of assays containing *Ec*MAT, QCMT and *Ec*MTAN showed full conversion of the peptide substrate (m = 2390.1 Da) after 20 h, indicated by a mass shift of +14 Da (m = 2404.1 Da). **b)** SAM regeneration assays with a 10× excess of substrate over AMP corresponding to a theoretical TTN of 20. Nearly full conversion of the peptide substrate to the methylated product was observed. **c)** Analysis of QCMT byproducts in SAM analogue assays containing PC*Mj*MAT and QCMT. L-Met was replaced with the respective analogue for *in situ* SAM analogue synthesis. All samples were taken after 20 h. While SAH (or the seleno-analogue) was observed in all assays, DOA was only detected when the substrate peptide was alkylated. **d)** Assays with PC*Mj*MAT, QCMT and *Ec*MTAN and L-Eth showed ethylation of the peptide substrate, indicated by a mass shift of +28 Da. **e)** Proposed reactions explaining the presence of SAH in samples containing *S*-allyl-D,L-homocysteine without detection of allylcobalamine. In the upper part, the *S*-allyl bond is cleaved rather than the *S*-C5' bond by QCMT, forming SAH and an allyl radical. In the lower part, the allyl residue is transferred to cobalamine by QCMT. The formed allylcobalamine rapidly degrades under formation of cob(II)alamine and an allyl radical.

### SAM regeneration from MTA supports aminopropyl transfer in polyamine synthesis.

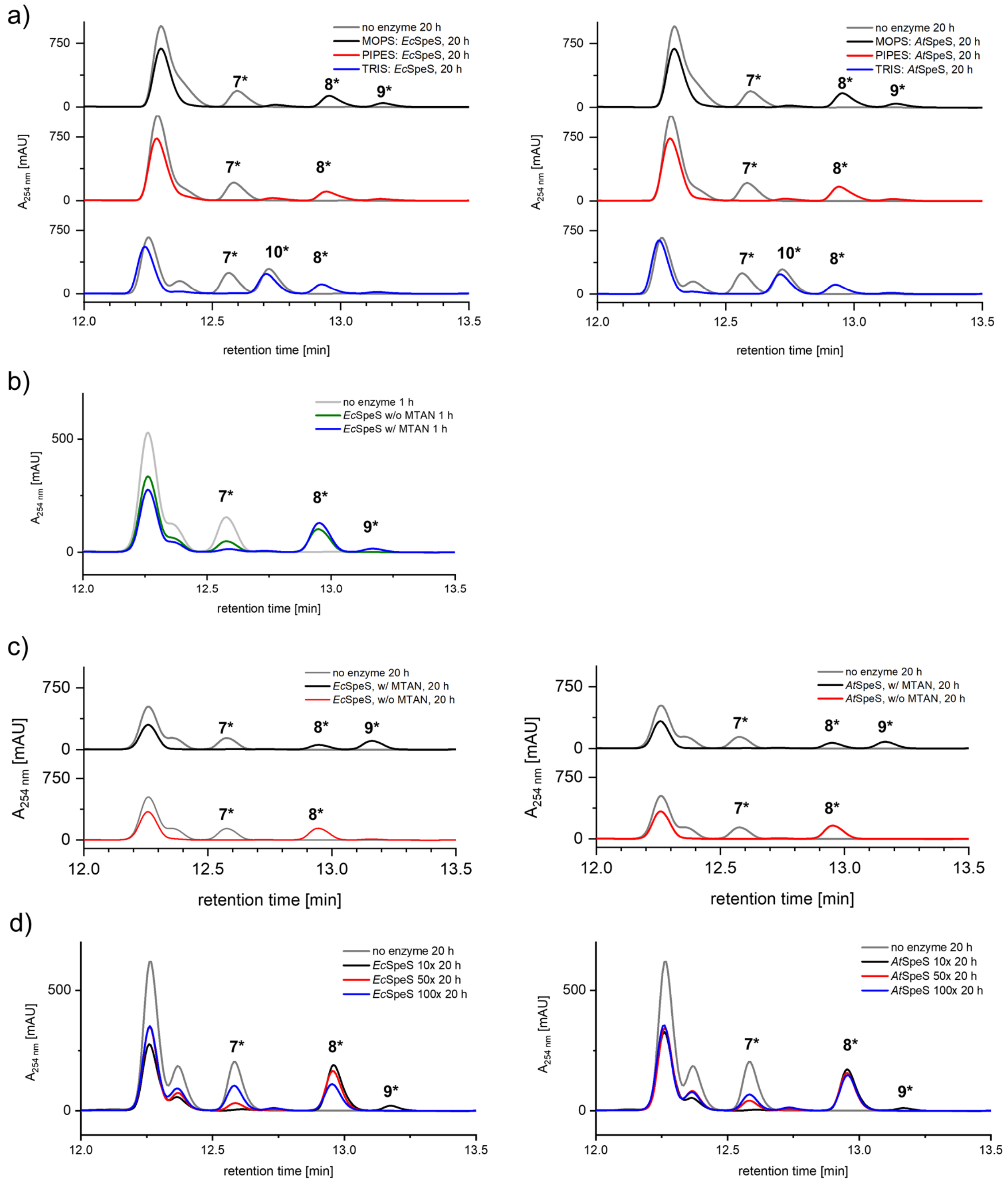

**Figure S5** Chromatograms of spermidine synthase assays. All samples were derivatised with BzCl (*) prior to analysis. **a)** Comparison of SpeS performance in the enzyme cascade consisting of *Ec*MAT, *Tm*SAMDc, SpeS and *Ec*MTAN. Tris buffer was compared to the tertiary amines MOPS and HEPES. In MOPS buffer most spermine (**9**) could be detected for *Ec*SpeS and *At*SpeS, indicating the highest activity. Note that Tris (**10**) interferes with analysis of putrescine (**7**) due to derivatisation with BzCl. **b)** *Ec*SpeS performance after 1 h in the linear cascade consisting of *Ec*MAT, *Tm*SAMDc, *Ec*SpeS with (blue) or without (green) *Ec*MTAN. Putrescine (**7**) is converted faster to spermidine (**8**) and spermine (**9**) when *Ec*MTAN is present. **c)** Assays with *Ec*MAT, *Tm*SAMDc and *Ec*SpeS or *At*SpeS were conducted with (black) and without (red) *Ec*MTAN after 20 h. For both SpeS, only small amounts of spermine (**9**) was observed without *Ec*MTAN, while assays with *Ec*MTAN led to production of a mixture of spermidine (**8**) and spermine (**9**). **d)** Typical chromatograms of 20 h SAM regeneration samples for *Ec*SpeS (left) and *At*SpeS (right) with either 10× (black), 50× (red) or 100× (blue) excess of putrescine (**7**) over AMP.

### Synthesis of L-methionine analogues.

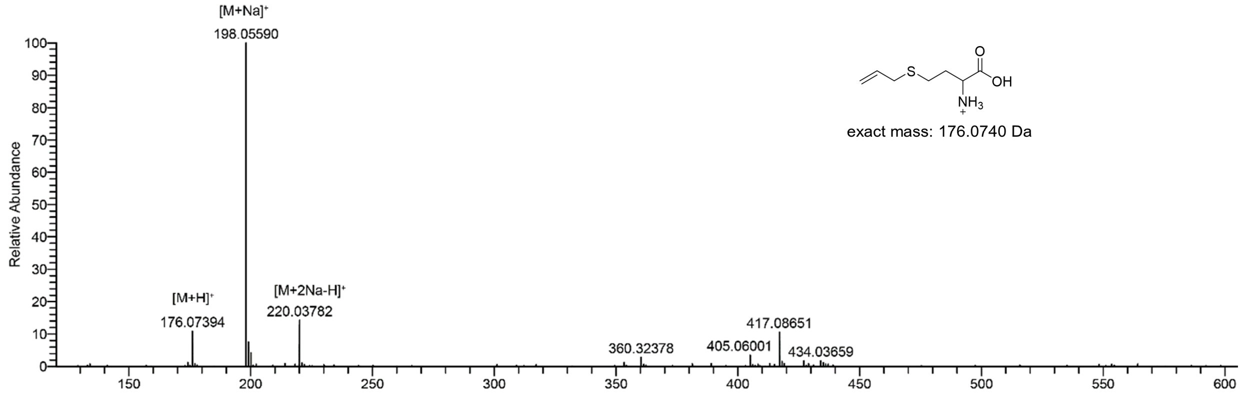

**Figure S6** HRMS (LTQ Orbitrap XL, Thermo Fisher Scientific) of allyl-D,L-homocysteine. Calculated for [M+H]+ = 76.07398 Da, found 176.07394 m/z.

a)

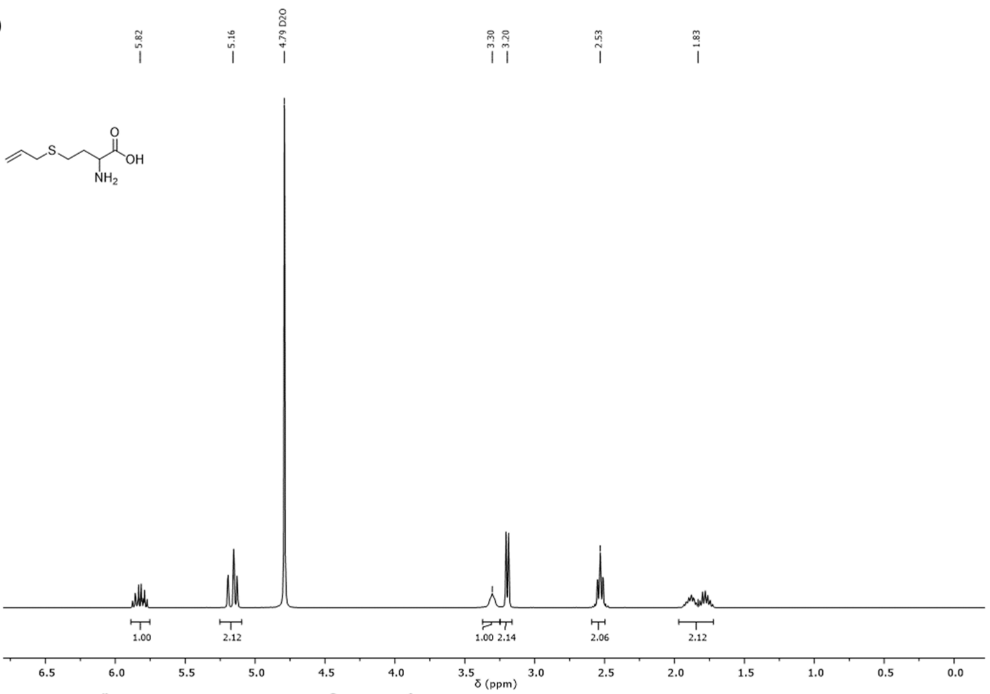

b)

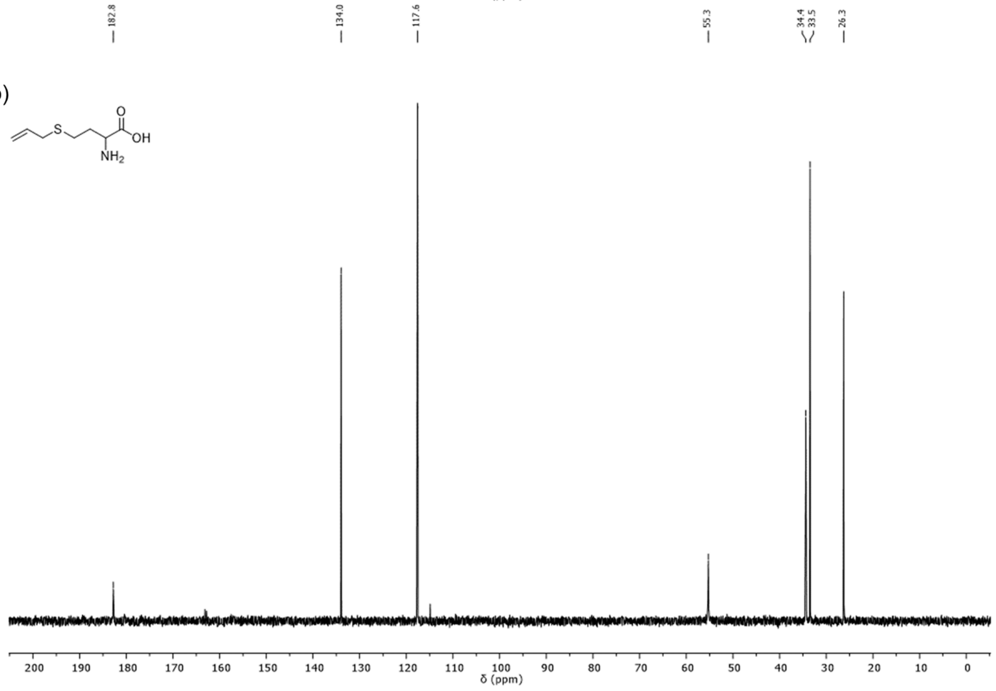

**Figure S7 a)** $^{1}$H NMR of allyl-D,L-homocysteine (400 MHz, $D_2O$ + 1 % NaOD) δ (ppm) = 5.82 (m, 1H), 5.31 – 5.10 (m, 2H), 3.30 (m, 1H), 3.20 (m, 2H), 2.53 (m, 2H), 1.98 – 1.67 (m, 2H). **b)** $^{13}$C-{1H}-NMR of allyl-D,L-homocysteine (101 MHz, $D_2O$ + 1 % NaOD) δ (ppm) = 182.8, 134.0, 117.6, 55.3, 34.4, 33.5, 26.3.

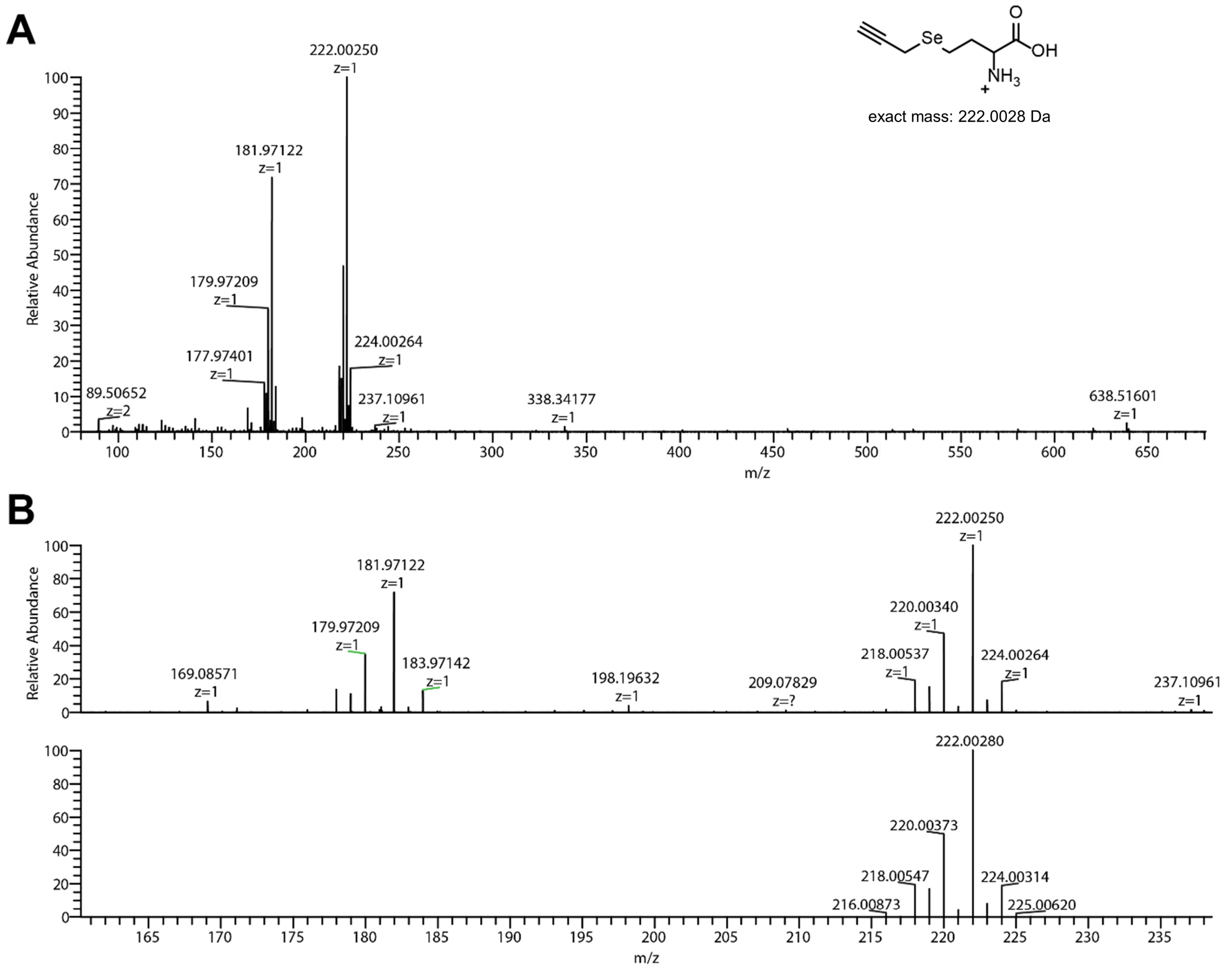

**Figure S8** HRMS of propargyl-L-selenohomocysteine. Calculated for [M+H]+ = 222.0028 Da, found 222.0025 m/z.

a)

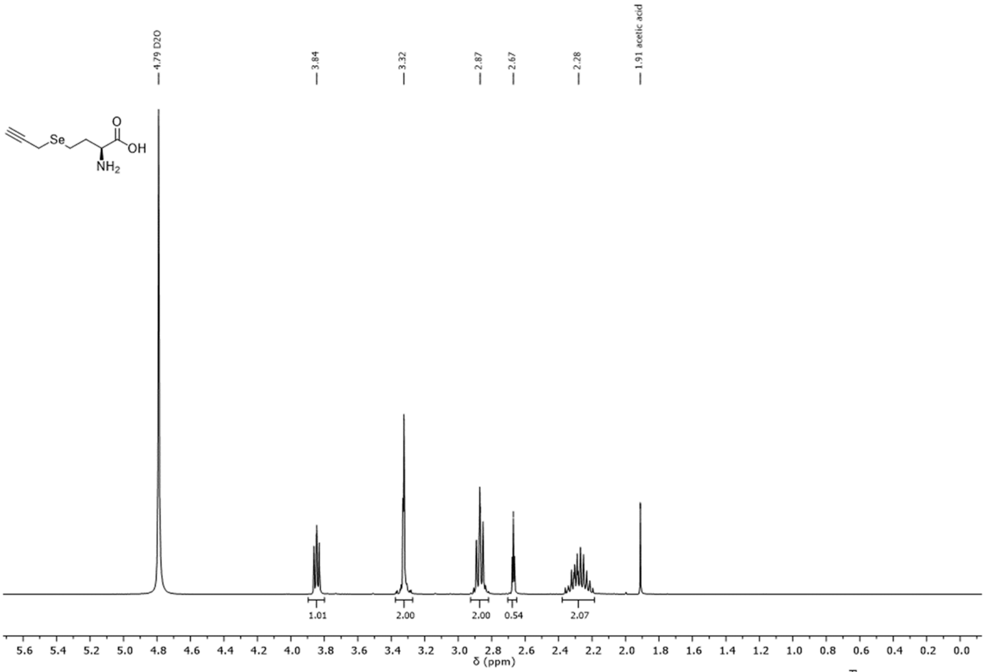

b)

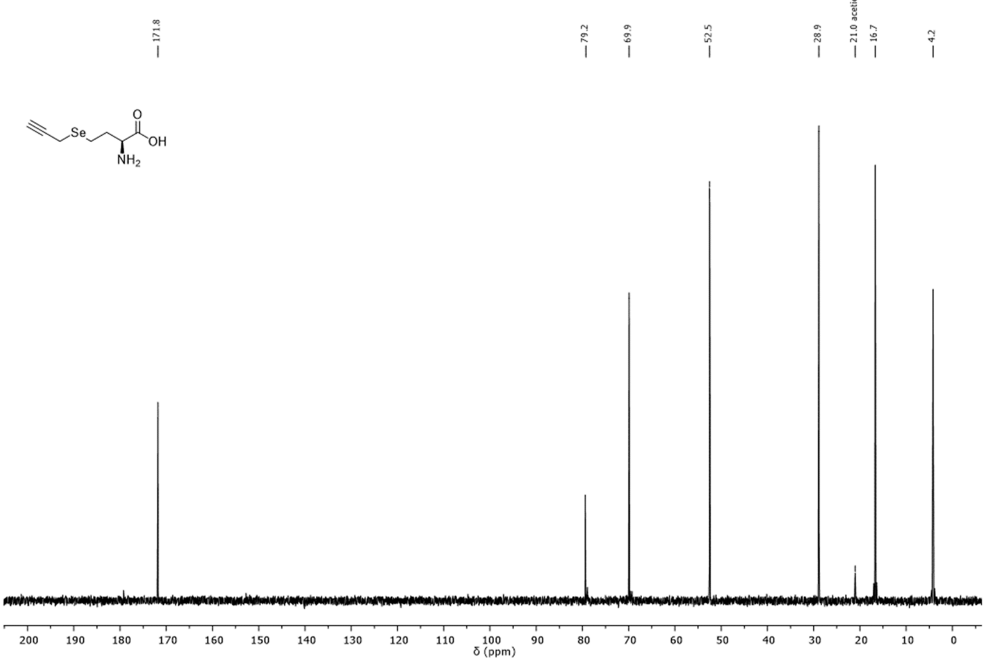

**Figure S9 a)** $^{1}H$ NMR of propargyl-L-selenohomocysteine (400-MHz, $D_2O$). δ (ppm) = 3.84 (t, $J$ = 6.9, 5.6 Hz, 1H), 3.39 – 3.28 (m, 2H), 2.92 – 2.82 (m, 2H), 2.67 (t, $J$ = 2.7 Hz, 1H), 2.38 – 2.19 (m, 2H). **b)** $^{13}C$-{$^{1}H$}-NMR of propargyl-L-selenohomocysteine (101 MHz, $D_2O$). δ (ppm) = 171.8, 79.2, 69.9, 52.5, 28.9, 16.7, 4.2.

### Author Contributions

JNA conceived and designed the study, supported by LG, LK and SM. LG, DM, and LK planned and performed experiments regarding ribose phosphorylation, cofactor supply and regeneration with MTs and radical SAM enzymes, supported by JNA, JG and GL. MK and MKFM planned and performed experiments regarding APTs, supported by JNA, XW and FPS. NVC and AR supported experiments with SAM analogues. LG, DM, LK, SJ, MK, and MKFM analysed the data. LG, DM, LK, JNA and CL interpreted the data, supported by MK, MKFM,

SJ, and HJJ. JNA, LG, and DM wrote the manuscript, supported by SM, HJJ, AR, GL, FPS, and CL. All authors reviewed and approved the final manuscript.
